# Nonspecific Membrane-Matrix Interactions Influence Diffusivity of Lipid Vesicles in Hydrogels

**DOI:** 10.1101/2023.02.03.526937

**Authors:** Nicky W. Tam, Otto Schullian, Amaia Cipitria, Rumiana Dimova

## Abstract

The diffusion of extracellular vesicles and liposomes *in vivo* is affected by different tissue environmental conditions and is of great interest in the development of liposome-based therapeutics and drug-delivery systems. Here, we use a bottom-up biomimetic approach to better isolate and study steric and electrostatic interactions and their influence on the diffusivity of synthetic large unilamellar vesicles in hydrogel environments. Single-particle tracking of these extracellular vesicle-like particles in agarose hydrogels as an extracellular matrix model shows that membrane deformability and surface charge affect the hydrogel pore spaces that vesicles have access to, which determines overall diffusivity. Moreover, we show that passivation of vesicles with PEGylated lipids, as often used in drug delivery systems enhances diffusivity, but that this effect cannot be fully explained with electrostatic interactions alone. Finally, we compare our experimental findings with existing computational and theoretical work in the field to help explain the non-specific interactions between diffusing particles and gel matrix environments.

**Statement of Significance:** The diffusion of nanoparticles in human tissues is dependent on interactions with the surrounding environment. This has wide implications for the development of nanoparticle-based therapeutics and drug delivery systems. Studying these interactions in human tissues and even in model hydrogels composed of reconstituted tissue components can be hampered by the many different complex interactions that can occur. By using a bio-inert hydrogel material like agarose, we remove the influence of specific biochemical interactions, allowing the study of how particle diffusion can be tuned with simple material properties like charge and rigidity. Taking advantage of these non-specific interactions, nanoparticles could one day be engineered to target specific organs by optimizing diffusion in certain tissue environments or retention and immobilization in others.

## Introduction

Large unilamellar vesicles (LUVs), or liposomes, are phospholipid structures 100-1000 nm in diameter that are often used as a minimal model of cell-derived extracellular vesicles (EVs). Their application in drug delivery systems takes advantage of the structure and function of their *in vivo* counterparts, as facilitators of intercellular transport (1, 2) to shield their payloads from the external tissue environment and mediate their transport to and uptake by target cells (3, 4). Despite the rising interest in using such lipid nanoparticle systems, for example to deliver anti-cancer therapeutics (5, 6) or as carriers of immunogenic materials in vaccine formulations (7), there is little data on how nanoparticle mobility and transport within tissues is affected by different tissue environmental conditions and the membrane material properties.

A 2009 study by Lieleg et al. (8) showed that hydrogel materials derived from extracellular matrix (ECM) can act as an electrostatic filter, sequestering charged nanoparticles while allowing neutral particles to pass through unimpeded. Later, Yu et al. (9) and Lenzini et al. (10) found that the deformability of lipid vesicles, modulated by the lipid composition of the membrane and by the presence of water-permissive channel proteins, respectively, can influence their access through hydrogel pore spaces and thus their movement and transport. Other work, including studies on rigid polymeric nanoparticles diffusing in polymer solutions (11, 12), in colloidal mucin suspensions (13), and in hydrogels (14–17) have also investigated the various ways in which charge and steric interactions affect the dynamics of non-deforming particles. Clearly, particle diffusion is affected by a diverse range of biophysical factors and of particular interest is the way these factors might interact. For example, recent theoretical work suggests that particle diffusability is a balancing act between particle deformability and particle-matrix adhesion.(18) When particles or vesicles are subjected to surface modifications, such as PEGylation (8, 13, 19–23) or the inclusion of more complex molecules, their surface interactions with the surrounding medium must also be taken into account. It is not difficult to imagine, then, that the combined effects of such interactions can give rise the complex distribution patterns of vesicles observed in vivo (24).

To systematically study how different material properties of lipid vesicles and ECM-like hydrogels can influence vesicle diffusion, we use single-particle tracking (25) to study the diffusion of synthetic LUVs embedded in agarose. Agarose is a polysaccharide polymer from red algae that undergoes thermo-reversible gelation via non-covalent hydrogen bonding (26, 27). While much simpler in chemical composition than the diverse molecules found in human ECM, agarose provides greater control over material properties. Stiffness and porosity of agarose gels, for example, can be found in a comparable range to human tissues such as brain or cartilage, and can be controlled with concentration and gelation conditions (28–30). Agarose is also a relevant material used in a number of different biomedical applications (28, 31, 32), including in three-dimensional cell culture platforms (33, 34) and as components of composite materials for tissue engineering (28, 35–37). Most importantly, agarose is bio-inert (33), allowing for the investigation of non-specific steric and electrostatic interactions without the influence of specific biochemical interactions that may be present with reconstituted ECM materials or mucin suspensions. By using this biomimetic system, we aim to better understand the biophysical mechanisms that govern the diffusion of extracellular-like vesicles or drug carriers in tissue-like materials. We also aim to directly compare deformable vesicles with similarly-sized rigid nanoparticles and particles with surface modifications to better tease apart how particle deformability and surface interactions contribute to overall particle diffusion and dynamics. Altogether, these results could one day lead to more efficient targeting and delivery of lipid nanoparticle-based therapeutics and vaccine delivery (3, 5–7, 38).

## Materials and Methods

### LUV production and characterization

Lipid stocks dissolved in chloroform (Avanti Polar Lipids, Alabaster, AL, USA) were used to prepare mixtures containing 4mM DOPC (1,2-dioleoyl-sn-glycero-3-phosphocholine) as a base solution. Negatively charged LUVs were made with a 2:1 molar ratio mixture of DOPC and DOPS (1,2-dioleoyl-sn-glycero-3-phospho-L-serine), while positively charged LUVs were made with a 2:1 molar ratio mixture of DOPC and DOTAP (1,2-dioleoyl-3-trimethylammonium-propane). PEGylated lipids were used to passivate vesicles for diffusion in hydrogels. At room temperature, all lipids are above their main phase transition temperatures and no demixing in the membrane is expected to occur. For PEGylated LUVs, additions of 1mol% or 10mol% DSPE-mPEG1K, DSPE-mPEG2K, or DSPE-mPEG5K (1,2-distearoyl-snglycero-3-phosphoethanolamine-N-[methoxy(polyethylene glycol)-1000], -2000], and -5000], respectively) were added to base solutions of DOPC or DOPC/DOTAP. Fluorescent visualization was facilitated by the addition of 0.2mol% DiIC_18_(5) (Thermo Fisher Scientific, Waltham, MA, USA; 1,1’-Dioctadecyl-3,3,3’,3’-Tetramethylindodicarbocyanine, 4-Chlorobenzene-sulfonate Salt).

LUVs were produced by first spreading a thin layer of a lipid mixture inside a glass vial and drying under vacuum for 1.5 hours. Next, the lipid film was hydrated with phosphate buffered saline (PBS; tablets for 200mL solutions from Sigma-Aldrich, St Louis, MO, USA) and vortexed for 30 minutes to produce multilamellar lipid structures. The resulting solution was then extruded with a Mini Extruder (Avanti Polar Lipids, Alabaster, AL, USA), 21 passes each through a 200nm and 100nm polycarbonate Nuclepore Track-Etched Membrane (Sigma-Aldrich, St Louis, MO, USA).

Size distribution and zeta-potential of particles were measured using a Malvern Instruments Nano-ZS Zetasizer equipped with a 632.8nm 4mW HeNe laser to ensure sample consistency. Samples in disposable folded capillary cells (DTS1070; Malvern Panalytical, Malvern, UK) were analyzed with dynamic light scattering (DLS) at a scattering angle of 173° to determine size distribution before determination of zeta-potential. All particles were measured in high-salt buffer conditions resulting in electrostatic screening, so zeta-potential values are used to illustrate relative differences in surface charge rather than absolute charge.

### Fluctuation analysis

To probe how the bending rigidity of lipid membranes changes with the presence of PEGylated lipids, we used fluctuation analysis on giant unilamellar vesicles (GUVs) (39, 40). GUVs were made using the gel-assisted swelling method (41, 42) (see Section S1 in the SI). Briefly, 20 µL 5% w/v solution of polyvinyl alcohol (PVA; fully hydrolyzed, MW = 145000 Da; Merck Group, Darmstadt, Germany) in water with 50 mM sucrose was spread onto a 2 cm by 5 cm area corresponding to the dimensions of a rectangular, 2 mm-thick Teflon spacer and allowed to dry completely in an oven at 50°C. Next, a thin 15µL layer of 4mM lipid mixture dissolved in chloroform was spread on top of the PVA layer and dried in a vacuum for 1.5 hours. The slide was then assembled into a sandwich with another glass slide and a Teflon spacer in the middle, held together with binder clips (Fig. S1). The lipid layer was hydrated for 30 minutes with 2mL PBS + 50mM sucrose (345mOsm/kg). The sucrose was necessary to help with the swelling process and to generate a sugar gradient that would later aid in visualizing the GUVs. GUVs were harvested and diluted 1:1 in a solution of PBS + 100mM glucose (394mOsm/kg) to slightly deflate the GUVs and were visualized under phase contrast with a 40× objective on a Zeiss AXIO Observer.D1 microscope. Image sequences of 3000 frames were recorded with a pco.edge sCMOS camera (Excelitas Technologies, Waltham, MA, USA) at 25 frames per second (fps) with 200µs exposure. Fluctuation analysis software (40) computed the bending rigidity based on the Fourier decomposition of thermally-driven membrane fluctuations into spherical modes. Fluctuation analysis as well as all other experiments were conducted at room temperature, approximately 23°C.

### Preparation and characterization of agarose gels

Stock solutions of 2% w/v low gelling-temperature agarose (BioReagent, for molecular biology; Sigma-Aldrich, St Louis, MO, USA) were made by dispersing agarose powder in PBS and microwaving at 350W power in 5-8s intervals until dissolved. Stocks were stored at 4°C and could be re-melted at 95°C using the same microwaving method. The molten agarose remained liquid down to 35°C. Gels of 1%, 0.5%, and 0.2% concentration were formed by melting stock gels and mixing with warm PBS (35°C) directly on glass slides for imaging, kept warm on a hotplate set to 35°C, or directly on a heated rheometer stage in the case of rheology measurements. Molten gels were taken off heating apparatus to cool to ambient temperature to induce gelation. Gel osmolality, which influences degree of vesicle deflation, was varied with the addition of glucose as opposed to salts to maintain the ionic strength of the solution, avoiding electrostatic screening effects. Solution osmolality prior to the addition of agarose was adjusted with a freezing-point osmometer (Osmomat 3000, Gonotec, Berlin, Germany). A list of tested gel formulations can be found in Table 2 in the SI (Section S5).

Bulk rheology of agarose hydrogels was studied in shear mode using an Anton-Paar MCR301 rheometer with 12mm cone-plate (CP12) geometry (Anton Paar GmbH, Graz, Austria). Gels were mixed directly on the rheometer stage heated to 35°C, then cooled below 20°C to allow the sample to start to set while the probe was lowered to the measurement position on the sample. The gel was left for 5 minutes to fully set before testing up to 1% rotational strain from 1-10Hz.

Average gel pore size was estimated using a turbidimetric assay described by Aymard et al. (29) and Narayanan et al. (27) Briefly, molten agarose was added to disposable 2.5mL PMMA cuvettes (Sigma-Aldrich, St Louis, MO, USA) and allowed to cool to ambient temperature (∼22°C) to gel. Absorbance values over 600-900nm were measured using a Thermo Spectronic Helios Gamma UV-Vis Spectrophotometer (Thermo Fisher Scientific, Waltham, MA, USA). This was compared to analytical data from Aymard et al. (29) (see Section S2, SI).

### Quantifying particle mobility

LUVs were embedded in agarose gels by mixing extruded LUV solutions with molten agarose directly on a glass microscopy slide within a rubber spacer (see Section S3, SI). A glass coverslip was placed on top, such that the agarose droplet wetted both glass surfaces, forming a disk. The imaging chamber was set aside at room temperature for 5 minutes to set. For control experiments with embedded polystyrene beads, working mixtures of Fluoresbrite YG 0.1µm-diameter Microbeads (Polysciences, Inc, Warrington, PA, USA) and FluoSpheres carboxylate-modified 0.1µm-diameter red (580/605) polystyrene beads (Invitrogen, Waltham, MA, USA) were made by diluting bead suspensions 1:100 in PBS before being mixed into gels, replacing the LUV solution at the same volume.

Samples were imaged with a pco.edge sCMOS camera mounted to a Zeiss AXIO Observer.D1 microscope with a 63× water immersion objective (Carl Zeiss AG, Oberkochen, Germany) in epifluorescence mode with appropriate excitation and emission filters. Image sequences of length 5s (∼100 frames) were captured with 20fps frame rate and ∼45ms exposure in a 100µm×100µm region of interest (ROI). Three ROIs were recorded per sample to account for internal heterogeneity. Particle mobility within gels was analyzed with the single-particle tracking plugin for FIJI developed by Sbalzarini and Koumoutsakos (25). A sequence length of 100 frames was chosen because longer sequences resulted in decreased signal-to-noise ratio from photobleaching, leading to increased false positives in particle detection. Histograms of the log_10_ diffusion coefficients (in m^2^s^-1^) obtained from the plugin were used to determine the mobile fraction.

Due to their size being below the diffraction limit, fluorescently-labelled particles with a nominal diameter of 100nm appear in images with pixel size 100nm×100nm as small clusters of 3-4 pixels with approximately 1- to 2-pixel spread (see Section S3, SI). In order for motion to be detected above the noise floor, a particle must be displaced more than 3 pixels from its original position, occurring over *t=*2.5s. This corresponds to the maximum lag time used in the calculation of the mean squared displacement (MSD), or half the total duration of the image sequences used. A theoretical lower limit of detection of particle movement can thus be calculated using the following relationship between the MSD and the diffusion coefficient, *D*, in two dimensions (43):

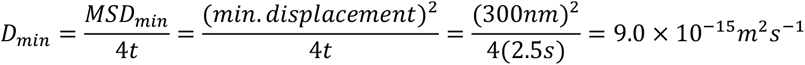

We thus use *log*_10_(*D*_*min*_) ≈ −14 as the cut-off point to determine whether a particle is mobile.

We also studied the infiltration of 100nm-extruded DOPC LUVs into preformed 1% w/v agarose hydrogels to obtain a collective diffusion coefficient. The prior imaging chamber setup was done with slight adjustments (see Fig. S4, SI). Briefly, agarose gel disks were formed without LUVs in an imaging chamber and allowed to set before a solution of LUVs (12µM lipid) was pipetted into the chamber. Three 100µm×100µm ROIs each in the gel interior and the exterior solution were imaged per time point, per sample over 100 hours. The number of particles in each ROI inside and outside the gel was counted as a function of time. Because of the slow diffusion and the low number of particles reaching the gel interior, we were unable to satisfactorily discretize the images to obtain smooth particle density gradients, as previously done for fluorescently labelled molecules (44). Instead, we obtained density gradients using finite differences and computed the diffusion coefficient as follows: each ROI is a rectangular box with dimension *h* = 100*µm* and *L* = 100*µm*. Assuming the depth of the observation volume is held constant, the system can be reduced to a two-dimensional model, whereby the two-dimensional flux per unit area, *J*, flowing into the ROI in the gel interior is given by

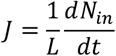

where *N*_*in*_ is the number of LUVs in the gel interior and *t* is time. Due to the slow diffusion, *N*_*in*_ appears to vary linearly in time (see Fig. S4D), hence

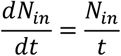

Fick’s first law of diffusion in two dimensions connects the density 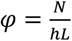 with the flux by introducing a diffusion constant, *D*, via

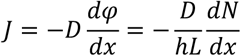

Finally, we approximate the density gradient using a finite difference

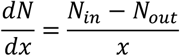

where *x* = 300µm is the distance between the ROI of the gel interior to the edge of the gel and *N*_*out*_ is the number of particles in the ROIs in the exterior LUV solution. Combining these relations and solving for the diffusion constant gives

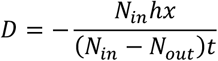

Because of the slow diffusion, all initial timepoints where *N*_*in*_ = 0 are excluded. Diffusion coefficients were calculated at each timepoint measured (4-13 data points per replicate over 100 hours of imaging, see Fig. S4D in SI), then averaged for each replicate.

### Statistical analysis

Histograms of log_10_ diffusion coefficients are normalized to show probability and represent pooled data from three ROIs within an individual gel. Each ROI corresponds to 100-300 diffusing particles. The variability in the number of identified particle tracks in different gels arises from differences in particle mobility. Statistics on mobile fractions were calculated with n=3 gels, presented as standard box plots showing the median (middle line), upper and lower quartiles (box limits), and full range of non-outlier data (whiskers). All other data are presented as mean with standard deviation. Statistical significance was determined with N-way ANOVA (as indicated) with Tukey-Kramer tests for multiple comparisons at the significance levels indicated, computed using MATLAB (MathWorks, Natick, MA, USA).

## Results and Discussion

### Mobility of embedded LUVs

LUVs embedded inside agarose gels were imaged with epifluorescence microscopy and analyzed with single-particle tracking (25) to obtain their diffusion coefficients (Fig. 1A,B). The mobile fraction was determined from the distribution of log_10_ diffusion coefficients. The peak observed at a value of -15 for 100nm-extruded LUVs composed of pure 1,2-dioleoyl-sn-glycero-3-phosphocholine (DOPC) embedded in 1% agarose (Fig. 1B) lies below the mobility cut-off of -14 (see Supplemental Information, SI Section S1) and thus corresponds to fully-immobilized particles. Particle immobilization and interactions in general with a gel matrix have previously been described in terms of electrostatic effects (at least for polystyrene particles) (45). In essence, while some particles can become fully entrapped by the gel matrix, other particles will be able to diffuse unhindered within the matrix voids as if they were in liquid water. Given close enough proximity to a wall or surface, particles can transiently bind and unbind with the gel matrix, reducing their mean squared displacement and thus “effective” diffusion coefficient. It is possible in our case with flexible lipid vesicles that these interactions can also be steric, with transient trapping and freeing of particles due to thermal fluctuations. The obtained effective diffusion coefficient can thus be used as a measure of the frequency and strength of membrane-matrix interactions, including steric ones. Since virtually all particles lie below -12 (Fig. 1B), the value given by the Stokes-Einstein equation for an ideal 100nm spherical particle diffusing in liquid water, this implies that all particles in the system are interacting with the gel matrix, sterically or otherwise.

**Figure 1.**
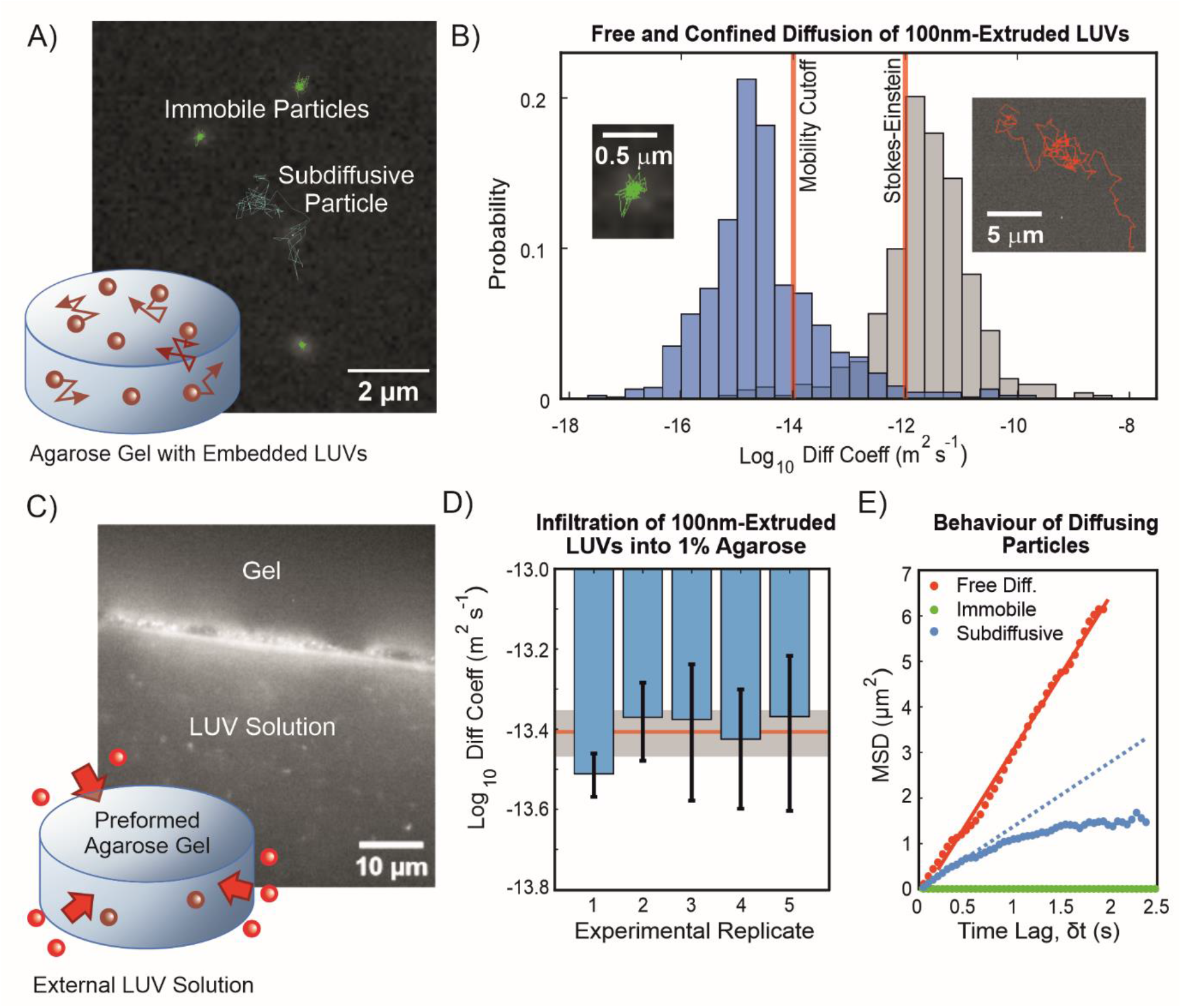
Particle mobility assessed in two ways: diffusion of DOPC LUVs embedded in gels and infiltration of LUVs into gels. A) The diffusion of 100nm-extruded DOPC LUVs embedded in 1% agarose gels is analyzed with single particle tracking. Particle paths are indicated as an overlay on a representative time frame. Here, green trajectories represent immobilized LUVs and a “subdiffusive” trajectory is shown in blue. This particular LUV was observed to “jump” between two hydrogel pore regions, resulting in this biphasic trajectory separated by a particularly large displacement. B) The base-ten logarithms of the diffusion coefficients are shown in histogram form for 100nm-extruded LUVs embedded in 1% agarose (blue) and in agarose-free liquid PBS (grey). The values of -14 and -12 are indicated by red vertical lines, the former being the lower limit of detection of particle motion for the experimental setup and the latter being the theoretical value determined from the Stokes-Einstein equation for an ideal 100nm-diameter spherical particle diffusing in liquid water. The insets show fragments of two trajectories corresponding to an immobilized/confined LUV in 1% agarose (green, left) and a freely diffusing one in agarose-free PBS (red, right). C) A measure of the collective particle diffusion is determined by incubating preformed agarose gel disks with an external solution of LUVs and monitoring their infiltration into the gel. The epifluorescence microscopy image shows the edge of an agarose gel disk, where LUVs can be seen adsorbed onto the surface. Individual LUVs in the solution are seen as tiny spots. D) The particle densities inside and outside the gel over time can be related to the diffusion coefficient (see SI Section S2). The average log_10_ diffusion coefficients computed from five independent experimental replicates are presented here with error bars showing standard deviation. The ensemble average across all experimental replicates was found to be 3.92±0.52×10-14m^2^s^-1^ (indicated as a red line with grey shaded area showing standard deviation), corresponding to a log_10_ value of -13.41. E) In both experiments, LUVs can be observed to exhibit different diffusive behaviours, characterized by different mean squared displacement (MSD) plots. A freely-diffusing LUV is characterized by a linear MSD plot, as shown in red, where the slope is proportional to the diffusion coefficient. The green plot shows a fully immobilized particle in agarose, where the slope is very close to zero. The blue MSD plot shows subdiffusive or anomalous diffusion behavior, whereby the MSD scales with a power of time, *MSD* ∝ *δt*^*r*^, *w*here r<1. A blue dotted line shows a linear fit of the first 10 data points of the MSD plot to illustrate the difference between subdiffusion and regular diffusion. The effective diffusion coefficient would be calculated and fitted over the whole MSD curve, resulting in an overall lower diffusion coefficient than would be expected of a freely diffusing particle.

Analyzing individual particle trajectories reveals different diffusive behaviours. Freely diffusing particles in liquid can be observed covering large areas while immobilized particles in gels remain stationary. Most LUVs embedded in gels undergo anomalous diffusion or subdiffusive behaviour, whereby particles diffuse within the confines of a matrix pore, resulting in a characteristic mean squared displacement (MSD) curve with a plateau at long time lags (Fig. 1E). These particles can sometimes “hop” between pore spaces, similar to what has been described in polystyrene nanoparticles diffusing in liquid polymer solutions (12) and in hydrogel matrices (14). Examples can be seen in Figs. 1 and 2. We note that particles appear to also become transiently trapped at certain locations within presumed pore spaces. These particles dwell at these locations for multiple consecutive image frames for periods of 0.25s, up to several seconds long, sometimes alternating between several trapping points before being freed. This appears to occur at size scales smaller than the apparent pore sizes mapped out by the rest of the particle’s trajectory or by neighbouring particles.

**Figure 2.**
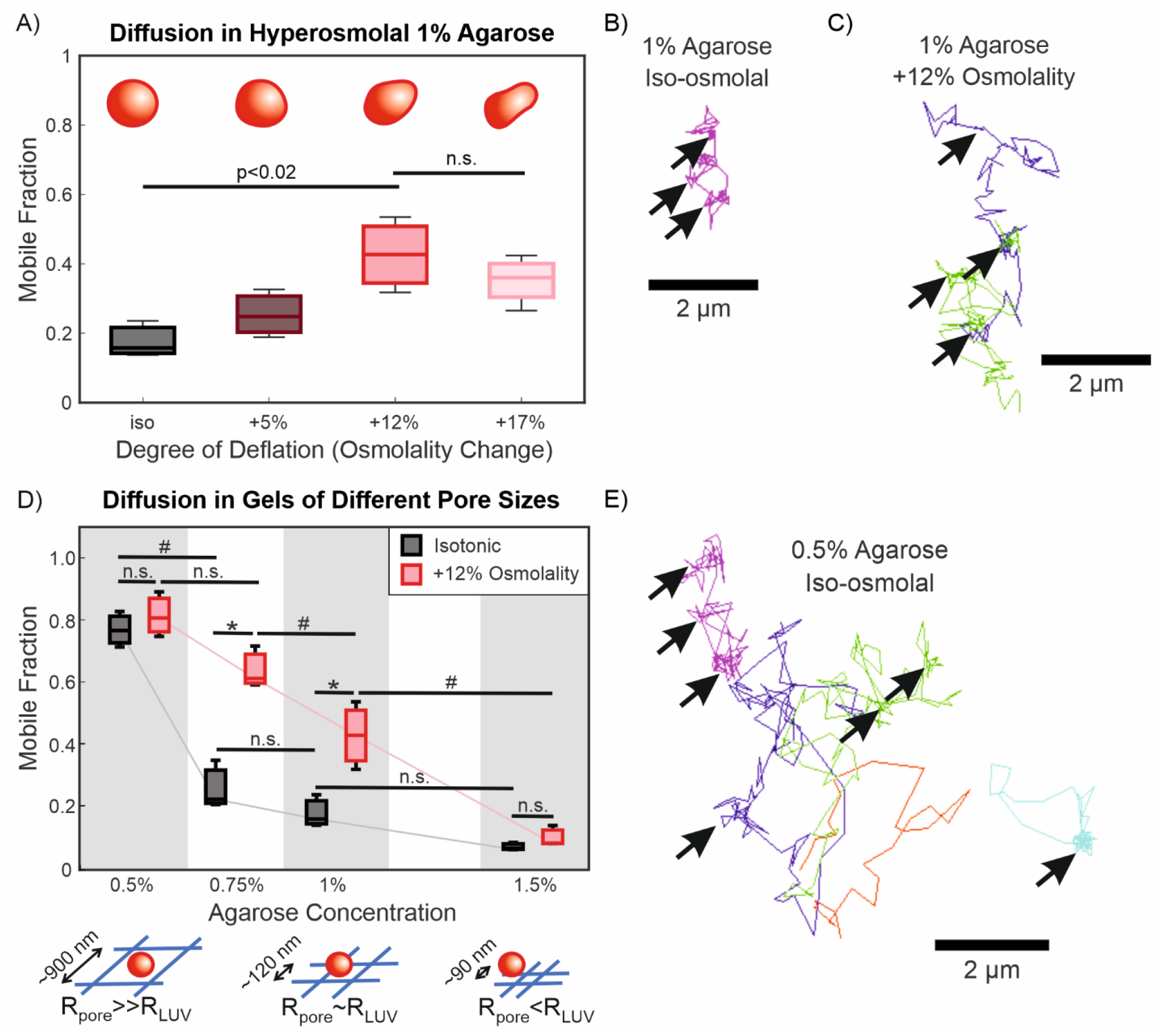
Mobility of DOPC LUVs increases with osmotic deflation and with hydrogel pore size. A) Mobile fractions of LUVs in agarose gels with different osmolalities relative to the LUV interior with cartoon representations of a LUV at varying stages of deflation. B) Example trace of a particle trajectory in 1% iso-osmolal agarose with arrows indicating apparent particle “trapping” regions. This trajectory may not be representative of the majority of particles in this condition, as it is specifically a particle with relatively high mobility to illustrate trapping behaviour, hence the apparent pore size traced by the particle trajectory may not correspond to the average size determined with turbidimetry. C) Representative particle trajectories in 1% agarose with 12% osmolality increase compared to the intravesicular solution. Different colours represent different particles and arrows indicate apparent trapping points. D) Mobile fractions of LUVs in agarose gels of differing concentration and osmolal strength with cartoon representations comparing the relative sizes of pores and LUVs. * represents a statistically significant difference between osmolalities at constant agarose concentration (p<0.01), while # represents a statistically significant difference between agarose concentrations at constant osmolality (p<0.05), as determined with one-way ANOVA and pairwise Tukey-Kramer post-hoc analysis. Lack of statistically significant difference (p>0.05) is indicated with n.s. E) Representative particle trajectories in 0.5% isoosmolal agarose with arrows indicating apparent trapping points. Different colours represent separate particles.

### Collective LUV diffusion from gel infiltration

We next looked at the ability of DOPC LUVs to infiltrate pre-formed agarose hydrogel disks to obtain an independent measure of particle mobility based on population dynamics (Fig. 1C, SI Section S2). While many LUVs ended up adhering to and getting stuck at the edge of the agarose disks, some LUVs were observed to infiltrate into them over 100 hours of imaging. Equilibrium in the density gradient was not reached in the time frame tested. By approximating the LUV density gradient with a finite difference, we calculate a population-wide diffusion coefficient using Fick’s First Law of Diffusion to be 3.92±0.52×10^-14^m^2^s^-1^ (see Fig. 1D and SI Section S4), corresponding to a log_10_ value of - 13.41. This falls between the mobility cut-off and the value for the Stokes-Einstein particle, agreeing well with the results from the single-particle tracking of embedded LUVs (Fig. 1A, B).

### Osmotic deflation of LUVs increases diffusivity

Work by Yu et al. (9) on LUV compositions of different phase transition temperature and by Lenzini et al. (10) on cell-derived EVs has shown that vesicle deformability can affect their diffusion in a hydrogel. Another way to make LUVs more deformable is to deflate them by introducing them into a hypertonic environment. We studied the mobility of DOPC LUVs in agarose gels of differing osmolalities by adding glucose to the hydrogel solution while keeping the initial intravesicular solution constant. Fig. 2A shows that LUV mobility increases with osmolality from isotonic to +12% osmolality, and thus degree of deflation. No significant difference in LUV size was detected with DLS, though size distributions appear to have slightly higher variability in hypertonic solutions (Fig. 3B in the main text; size distributions found in Fig. S5C in the SI). We also did not observe differences in the bulk rheology of agarose gels formed with and without glucose (Fig. S5A), thus the microstructure of the gel is not expected to vary more than what is naturally found in agarose (26). The lack of statistically significant difference in mobility from +12% to +17% osmolality (Fig. 2A) could be due to a phenomenon similar to what was described by Yu et al. (9) whereby greater deformability ultimately exposes greater surface area that can conform to and interact with the matrix walls, resulting in immobilization. Recent theoretical work (18) also shows that particle diffusibility in a gel matrix is dependent on a balance of particle deformation and adhesive forces in the matrix. Kinetic energy in a hyper-deformable vesicle’s collision with a matrix wall could end up being spent on deforming the membrane, such that insufficient energy remains for overcoming matrix-adhesive forces.

**Figure 3.**
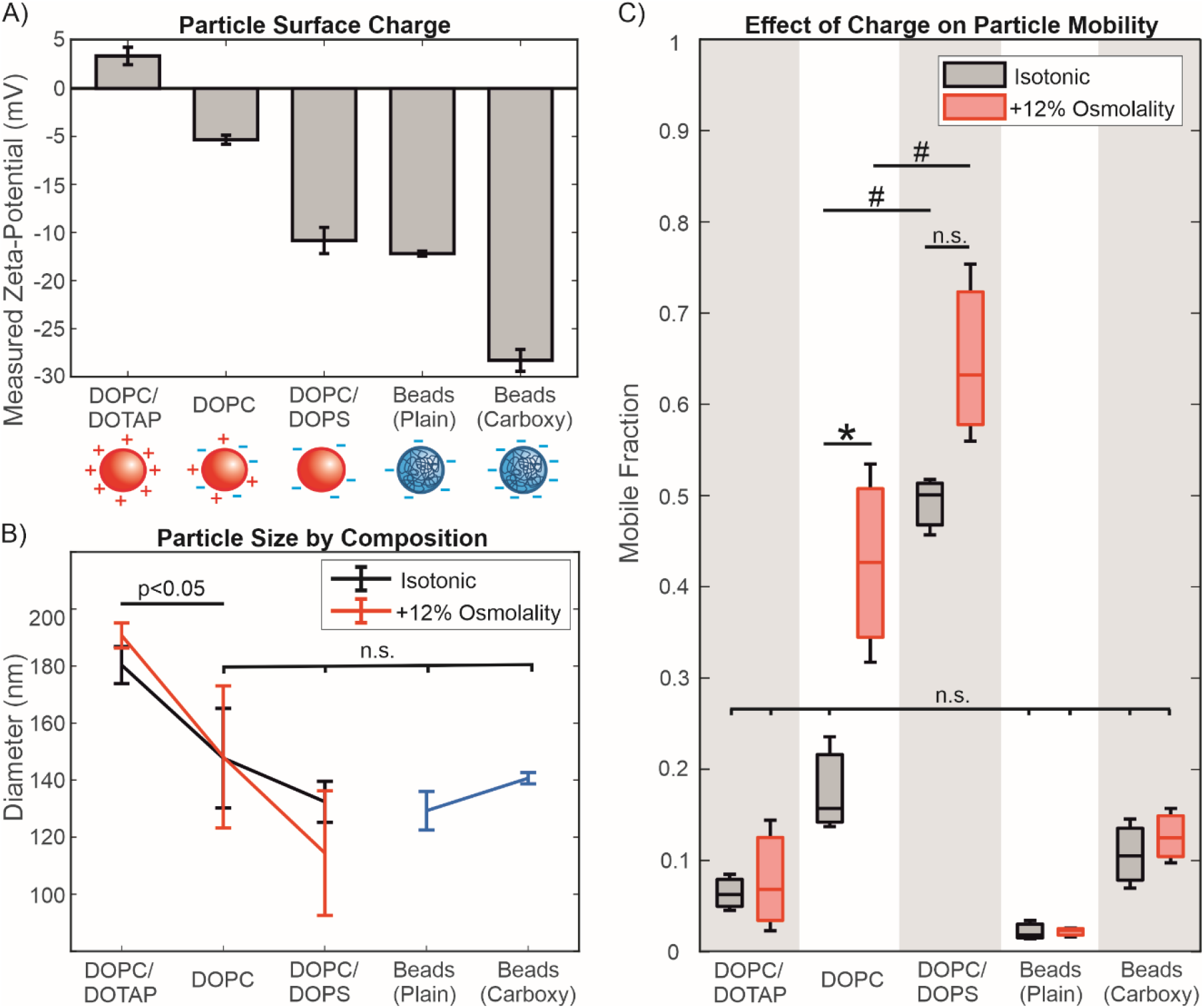
Effect of charge and composition on particle mobility in 1% agarose gels. The lipid ratios in DOPC/DOTAP and DOPC/DOPS LUVs correspond to 2:1. A) Zeta-potential of LUVs and polystyrene beads (either plain or with surface carboxylation), as measured in PBS; the sketches below roughly illustrate their surface charge. B) Diameters of LUVs and particles, as measured with dynamic light scattering. Average diameters of DOPC/DOPS and DOPC LUVs, as well as the polystyrene beads (blue) are not significantly different, in both isotonic (black) and hypertonic (red) conditions, as determined by 2-way ANOVA with pairwise Tukey-Kramer post-hoc analysis (p>0.05). Representative size distributions of particles can be found in Fig. S6 in the SI. Sizes of DOPC/DOTAP LUVs are significantly larger than those of the other particles (p<0.05). C) Mobile fractions of different particles in isotonic (black; 290mOsm/kg) and +12% hypertonic (red; 320mOsm/kg) buffer conditions; see Fig. S5 for diffusivity data. Statistical significance was determined with 2-way ANOVA with pairwise Tukey-Kramer post-hoc analysis. # represents a significant difference across membrane compositions (p<0.01). Statistically significant differences (p<0.02) at different osmolalities is represented with * while lack of a significant difference (p>0.02) is denoted with n.s. The lack of a statistically significant difference between DOPC/DOPS LUVs in isotonic and hypertonic environments was confirmed with a paired-sample t-test, which did not reject the null hypothesis (p>0.05). Note that although the maximum extents of the data (whiskers) do not overlap, the range of the 95% confidence intervals (not shown for clarity) do overlap.

When the pore size of the gel is increased from 120±6nm to 250±30nm by decreasing the agarose concentration from 1% to 0.75% w/v (Fig. S2), the mobility of LUVs does not increase significantly in isotonic conditions, but does so under hyperosmotic conditions (Fig. 2D). A further increase in pore size to 900±300nm by decreasing the agarose concentration to 0.5% results in an overall increase in LUV mobility. At this concentration of agarose, the difference between the mobile fractions in different osmolalities is not statistically significant. The effect of deflation thus appears to only be relevant when the average pore diameter is comparable (i.e., on the same order of magnitude) to the diameter of the LUV. This is reasonable, as the LUVs would have greater access to matrix pores regardless of deformability in the 0.5% gel. At the opposite extreme, decreasing the average pore size to below the average LUV diameter to 90±6nm results in nearly full immobilization of LUVs, even when osmotically deflated.

### LUV surface charge affects mobility

Lieleg et al. (8) reported that Matrigel, a complex mixture of cell-derived ECM materials exhibits electrostatic filtering behavior on diffusing particles. This has also been shown with polymer solutions and in computer simulations (11). To determine whether agarose has similar characteristics, we produced negatively charged LUVs from a 2:1 molar ratio mixture of DOPC/DOPS (1,2-dioleoyl-sn-glycero-3-phospho-L-serine ) as well as positively charged LUVs composed of 2:1 DOPC/DOTAP (1,2-dioleoyl-3-trimethylammonium-propane) to embed in agarose (Fig. 3A). The addition of DOPS does not significantly change LUV size (Fig. 3B), but the DOPC/DOTAP particles appear significantly larger than other tested particles, possibly due to aggregation. Figure 3C shows that the negatively charged DOPC/DOPS LUVs have greater mobility overall compared to the positively charged DOPC/DOTAP and pure zwitterionic DOPC LUVs (see also Fig. S6). Although there appears to be an increase in mobility in DOPC/DOPS LUVs upon deflation, this difference is not statistically significant according to a one-way ANOVA test with pairwise comparisons over the whole data set (p>0.05). This was also confirmed with a paired-sample t-test with just the DOPC/DOPS data, which did not reject the null hypothesis (p>0.05). DOPC/DOTAP LUVs remain immobile in hypertonic conditions, as well as in much lower agarose concentrations (Fig. 4C). At 0.5% w/v, agarose would have an average pore size of 900nm, which should be large enough to accommodate even the largest aggregates detected with DLS (∼300nm, see size distributions in Fig. S6B). At 0.2% w/v, agarose behaves like a liquid, being able to flow. The lack of mobility at these concentrations would suggest that this interaction is not merely steric, but electrostatic, causing the positively charged LUVs to stick to the agarose polymer bundles. This could help explain the apparent size of DOPC/DOTAP LUV aggregates seen in agarose gels (Fig. 4A), but not in suspension. Analysis of fluorescence intensity profiles of particles embedded in agarose shows that such aggregates approach 1µm in diameter, while the maximum particle size detected with DLS is approximately 300nm.

**Figure 4.**
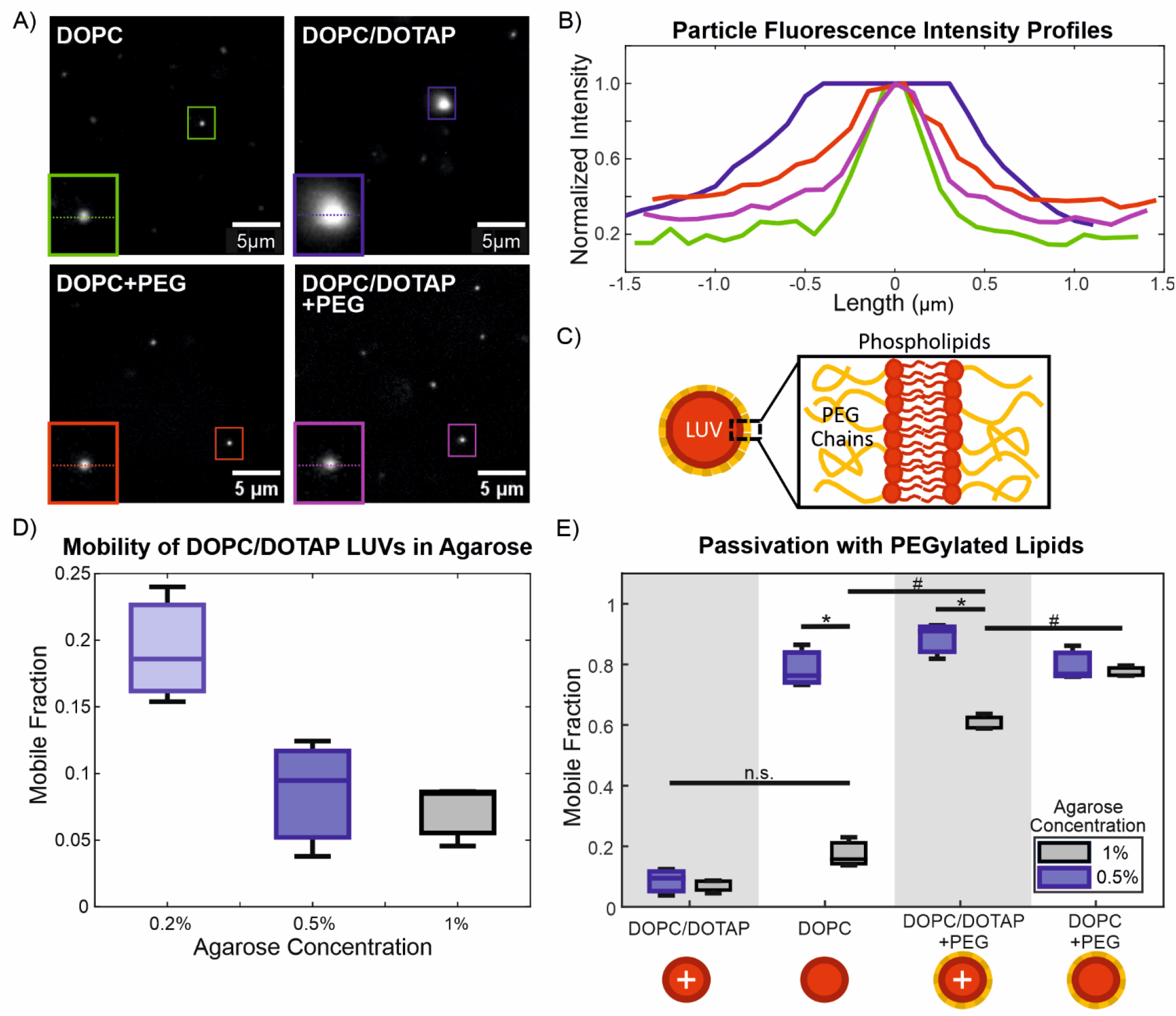
Recovery of DOPC/DOTAP LUV mobility with PEGylated lipids. A) Representative images of 100nm-extruded DOPC, 2:1 DOPC/DOTAP, DOPC +10mol% DSPE-mPEG1K (DOPC+PEG), and 2:1 DOPC/DOTAP +10mol% DSPE-mPEG1K (DOPC/DOTAP+PEG) LUVs in 0.5% agarose gels. Selected particles are indicated with coloured boxes and shown in insets (2.9µm width), where a dotted line indicates the position at which the fluorescence intensity profiles in (B) are taken. Particles appearing with different absolute intensities could be due to being at different focal depths or because of different degrees of photobleaching, as particles move in and out of frame or focus. The selected DOPC/DOTAP particle is likely an aggregate of LUVs. B) Normalized fluorescence intensity profiles of selected LUVs. Colours correspond to the particles in (A). The DOPC, DOPC+PEG, and DOPC/DOTAP+PEG profiles represent single particles, which are smaller than the diffraction limit of our imaging setup. These particles thus appear blurry with considerable spread, but with comparable normalized fluorescence intensity profiles. The DOPC/DOTAP aggregate, by contrast, has a diameter approaching 1µm and is noticeably larger by visual inspection. C) Schematic diagram of a LUV membrane containing PEGylated phospholipids. D) Mobile fractions of 2:1 DOPC/DOTAP LUVs in agarose gels/solutions of differing concentration. Agarose at 0.2% concentration behaves macroscopically as a liquid, but likely has some weak molecular network linkages. E) Comparison of mobile fractions of 2:1 DOPC/DOTAP LUVs (DOPC/DOTAP), pure DOPC LUVs (DOPC), 2:1 DOPC/DOTAP LUVs +10mol% DSPE-mPEG1K (DOPC/DOTAP + PEG), and DOPC LUVs +10mol% DSPE-mPEG1K (DOPC + PEG). # indicates statistically significant differences when comparing mobilities across LUV compositions in the same gel concentration. * indicates statistically significant differences when comparing mobilities of LUVs in different agarose gel concentrations. Mobility of DOPC/DOTAP LUVs in both 1% and 0.5% agarose, as well as that of DOPC LUVs in 1% agarose are not significantly different (n.s.). Statistical significance is determined with two-way ANOVA with pairwise Tukey-Kramer post-hoc analysis for multiple comparisons.

As a control, we compared LUV mobility to that of polystyrene beads. Despite having a similar size and negative surface charge to our DOPC/DOPS LUVs, plain polystyrene beads are fully immobilized in the hydrogel compared to the highly mobile DOPC/DOPS LUVs. Beads with surface carboxylation appear to have slightly higher mobility than plain beads, but not at a statistically significant level. Neither types of beads are affected by increased osmolarity. One explanation for this relates to the fact that the polystyrene beads are rigid while the LUVs are deformable and capable of squeezing through gel matrix pores that would otherwise be too small to pass through. The increased surface charge of the DOPC/DOPS compared to DOPC LUVs should also result in a slightly stiffer membrane (46), although this effect could be minimized by the high salt concentration. In 0.5% agarose gels, plain polystyrene beads remain immobile (mobile fraction = 0.07 ± 0.05) despite the much larger pore size while carboxylated beads become much more mobile (mobile fraction = 0.87 ± 0.08; see Fig. S6 in SI). It is possible that the enhanced negative charge of the carboxylated beads overcomes specific attractive interactions present between the agarose gel and the polystyrene beads. Another possibility relates back to the theoretical work of Yu et al. (18), who showed that highly rigid particles require very low attractive forces to maintain diffusibility. These results reveal an interesting intersection of different factors affecting the diffusion of particles through a gel matrix. In particular, we note that the use of polystyrene nanoparticles to model the diffusion of LUVs, EVs, and other soft particles may not be accurate due to the differences in their overall deformability, even if their surface properties are matched.

### PEGylation of LUVs increases their mobility

Nanoparticles are often “passivated” with PEG, a hydrophilic polymer used to prevent the adsorption of proteins on surfaces and hinder or slow down immune reactivity. It has been claimed that this passivation effect is due to the neutral charge of PEG masking the underlying surface (8, 19, 22), though other works suggest the involvement of steric or entropic effects of the PEG chains (13, 47). We questioned whether this effect could restore the mobility of our DOPC/DOTAP LUVs (Fig. 4C). For this to work, the layer of PEG chains would need to be thicker than the Debye length of the charges on the membrane. The inclusion DSPE-mPEG1K ((1,2-distearoyl-sn-glycero-3-phosphoethanolamine-N-[methoxy(polyethylene glycol)-1000]), a phospholipid coupled to a 1000Da PEG chain at 10mol% should make a PEG layer thick enough to screen out most electrostatic effects. In the high ionic strength buffer environment (PBS) that was tested, the Debye length, the distance over which an electric charge exerts an influence is <1nm.(48) Meanwhile, the three-dimensional Flory radius of the polymer, *R*_*F*_ is given by 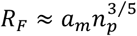, where *a*_*m*_ is the size of the monomer unit (*a*_*m*_ ≈ 0.39nm *for* PEG, as used by Marsh et al. (49)) and *n*_*p*_ is the number of monomers in the polymer (∼23 for PEG1000). The mean-field theory equilibrium length of the polymer chain, *L*^*MF*^, describing the average height of the polymer brush layer is also given by Marsh *et al*. as 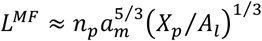 where *X*_*p*_ *is* the molar fraction of PEGylated lipid and *A*_*l*_ *is* the area per lipid molecule of the membrane, taken to be ∼0.6nm^2^ for a lipid in the fluid phase. Both *R*_*F*_ and *L*^*MF*^ are approximately 2.6nm for PEG1000 at 10mol% coverage in a fluid-phase lipid membrane, and thus greater than the Debye length. Therefore, the PEG layer should be thick enough to block electrostatic interactions with the underlying phospholipid surface. This indeed appears to restore the mobility of the DOPC/DOTAP LUVs in 0.5% w/v agarose to the same level as pure DOPC LUVs (and DOPC+PEG LUVs; Fig. 4D). The mobility of the passivated DOPC/DOTAP+PEG LUVs in 1% agarose is much improved compared to that of the non-passivated DOPC and DOPC/DOTAP LUVs, but is lower than the passivated DOPC+PEG LUVs. One possible explanation for this is that LUVs need to deform to fit through the smaller pores of the 1% agarose. This could force the PEG layer to compress or deform out of the way, exposing the underlying positive charge. This is possible, as previous work has shown that PEG is highly compliant (20, 23). Furthermore, the addition of PEG appears to prevent the particle aggregation apparent in DOPC/DOTAP LUVs, bringing the mean peak value of the size distribution closer to the expected value of 100nm (Fig. S6; SI). It also seems to prevent aggregation of particles upon embedding in agarose (Fig. 4A,B). Since free particles have a diameter below the diffraction limit of our imaging setup, we cannot directly compare their apparent sizes in images. Direct comparison of fluorescence intensity is also not possible due to the particles appearing at different focal depths with our epifluorescence imaging and because of photobleaching. However, DOPC/DOTAP LUV aggregates are apparent upon visual inspection and analysis of their fluorescence intensity profiles shows that they have diameters approaching 1µm. Aggregates of these sizes are not detected by DLS and are not apparent with visual inspection of samples in liquid suspension, only appearing upon embedding in agarose. These aggregates, thus, are likely the result of electrostatic interactions with the agarose matrix, causing the pinning together of particles. This effect disappears upon addition of PEGylated lipids, as the PEG prevents both particle-particle and particle-matrix interactions.

While PEGylation improved the mobility of DOPC/DOTAP LUVs, it also improved that of DOPC LUVs despite previous results of higher mobility in agarose with greater negative charge. DOPC LUVs also do not appear to aggregate as DOPC/DOTAP LUVs do, so this effect cannot be due to the prevention of particle aggregation. To investigate further how PEG affects LUV mobility, we tested other PEG chain sizes (1000, 2000, 5000 Da) at two concentrations (1mol% and 10mol%; Fig. 5A-C). Figure 5D,E shows that all PEG chain sizes improve LUV mobility in 1% agarose, while LUVs with PEG2000 and PEG5000 appear to be more mobile than those with PEG1000 in 0.5% agarose gels (see histograms and size distributions in Fig. S7, SI). LUV mobility is higher with 10mol% PEGylated lipid, where the PEG chains are in a polymer brush conformation (20, 23, 49) compared to 1mol% polymer, where they are in mushroom conformation (20, 23, 49). The difference in PEG chain conformations could help explain why PEG1000 appears ineffective in 0.5% agarose, as the mushroom-to-brush transition occurs at a higher concentration and is less well-defined. Regarding the distributions of the log_10_ diffusion coefficients (SI, Fig. 7A), although a greater proportion of particles become mobile upon addition of PEGylated lipids, the mean peak value of the mobile particles does not appear to change for a given concentration of agarose. For LUVs in 1% agarose, the peaks of the distributions fall consistently below -12 and appear to shift towards greater mobility, approaching -12 when the agarose concentration is decreased to 0.5%. This does not appear to be affected by size or surface coverage of PEG chain. While the PEG might prevent matrix interaction and immobilization, the agarose gel would continue to impose steric hindrance and restrict the space a particle can diffuse in, resulting in subdiffusion, and thus, a lower effective diffusion coefficient. When this steric hindrance is lessened by the much larger matrix pores of 0.5% agarose, particles are able to diffuse unrestricted over a much larger area, approaching the Stokes-Einstein-predicted value of -12 for a similarly sized particle in liquid.

**Figure 5.**
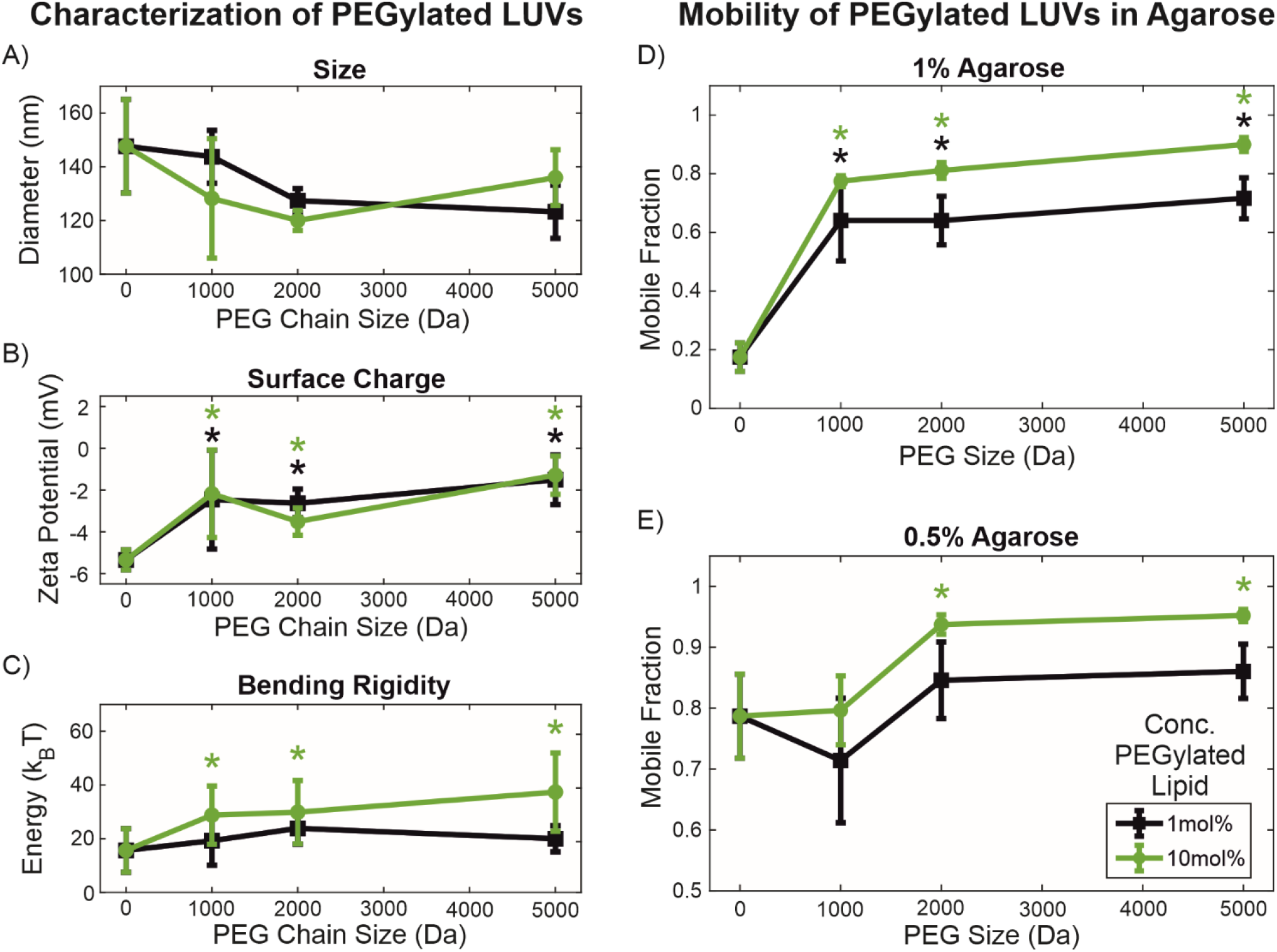
Effect of degree and density of PEGylation on LUV characteristics and mobility. Green data represents 10mol% concentration of PEGylated lipid in membranes while black data represents 1mol% concentration. A) Influence of PEG chain size and concentration on LUV size, as measured with DLS. No statistically significant differences were found (p>0.05) and size distributions appear similar (Fig. S7B in SI). B) Influence of PEG chain size and concentration on surface charge, measured as the zeta-potential. Bare DOPC LUVs are significantly more electronegative compared to PEGylated LUVs, regardless of PEG size and concentration (p<0.01, green and black *), demonstrating screening by PEG. C) Influence of PEG chain size and concentration on membrane bending rigidity, measured with fluctuation analysis on giant unilamellar vesicles (GUVs) and presented in k_B_T energy units (k_B_ being the Boltzmann constant and T being temperature). Membranes with 10mol% PEGylated lipid are significantly stiffer than those with 1mol% PEGylated lipid. Bare DOPC membranes are significantly less stiff than PEGylated membranes only at 10mol% concentration (p<0.01). No statistically significant differences were found between different PEG chain sizes. Statistically significant differences compared to bare DOPC membranes were determined with 2-way ANOVA with pairwise Tukey-Kramer post-hoc analysis, as indicated with *. D) Mobilities of all PEGylated particles in 1% agarose are significantly greater than those of bare DOPC LUVs at both concentrations of PEGylated lipid (p<0.01, green and black *). Mobilities of LUVs with 10mol% PEGylated lipid are significantly greater than those of LUVs with 1mol% PEGylated lipid (p<0.01). E) Mobilities of LUVs with 10mol% PEG2000 and PEG5000 are significantly greater than those of bare LUVs and LUVs with 10mol% PEG1000 (p<0.01). No significant differences were found at 1mol% PEGylated lipid. Statistically significant differences compared to bare DOPC membranes are indicated with *, as determined by 3-way ANOVA with pair-wise Tukey-Kramer post-hoc analysis.

These results suggest that the effect that PEG has on particle mobility is not entirely electrostatic. Greater mobility is also unlikely to be due to disaggregation of particles, since DOPC LUVs do not appear to aggregate without PEG upon visual inspection and according to the lack of change in average size and size distributions measured with DLS (Fig. 5A and Fig. S7B).

Fluctuation analysis on GUVs composed of the same lipid mixtures as the tested LUVs shows a slight stiffening of lipid membranes with the presence of PEGylated lipids (Fig. 5C). This behavior is consistent with predictions for membrane stiffening by anchored polymers,(50) and experiments on membranes with biopolymer adsorption(51) as well as microemulsions.(52) However, the stiffening observed here does not explain the increase in LUV mobility measured when PEGylated lipids are present. As demonstrated in Fig. 2, mobility should improve with overall deformability, which would decrease with increasing bending rigidity. One possibility would be that the PEG forms a soft lubricating layer that facilitates the movement of LUVs through matrix pores. This would be supported by the higher mobile fraction observed with larger PEG chains and at higher PEG coverage (in the brush regime), which would form a thicker layer. The underlying mechanism for this could be explained by the entropic repulsive force described in the computational work of Li and Shi,(47) whereby the compression of the PEG layer during a collision with the matrix wall would produce a strong repulsive force, preventing entrapment of the LUV in the matrix. It has also previously been reported that surface PEGylation improves the mobility of nanoparticles in mucin gels by sterically preventing the adsorption of colloidal mucin onto the particle surface.(13) These results, though similar, are likely due to a different mechanism, as agarose is not known to exist in a colloidal phase capable of adsorbing onto the particle surface and should be fully incorporated into the matrix scaffold.(26, 30) A particularly interesting question for future consideration would be whether the glycocalyx, the diverse array of polymeric sugar molecules expressed on the surfaces of cells and cell-derived EVs might play analogous roles in vivo.

## Conclusions

Here, we have demonstrated the applicability of agarose as a bio-inert and non-adhesive model for investigating the non-specific steric and electrostatic interactions of vesicles with a 3D polymer matrix. The use of a bottom-up biomimetic approach to study lipid vesicle diffusion in hydrogel materials has shown that different biophysical factors contribute to particle dynamics. Even without specific biochemical interactions via cell adhesion molecules and ECM proteins, the combined effects of non-specific steric and electrostatic interactions can give rise to selective filtering behavior, allowing particles of certain surface charge characteristics and overall deformability to diffuse freely, while entrapping and immobilizing others with different propertiees. In vivo, such preferential infiltration of certain vesicle populations into tissues with specific ECM composition or architecture may potentially form a basis for organotropic “homing” of particles.

We have presented experimental data that support and bring together existing computational and theoretical work on particle diffusion through different ECM-like environments. We show that polystyrene beads are a poor model for studying the diffusion of liposomes and that the effect of PEGylation on particle diffusivity is not due to electrostatic effects, as previously claimed, nor through the prevention of adsorption of soluble or colloidal materials on the particle surface, as no such species exist in our system. The design of lipid nanocarriers and engineered liposome-based therapeutics can thus take advantage of non-specific membrane-matrix interactions to achieve improved penetration and retention in target tissues. Further research on ECM-derived materials and more complex lipid vesicle systems is needed to better understand the diverse molecular interactions that govern the movement of vesicles in the tissue microenvironment.

## Acknowledgements

N.W. Tam would like to acknowledge funding from the International Max Planck Research School on Multi-Scale Biosystems, as well as Alexander Becker, a summer intern supported by the German Academic Exchange Service (DAAD) RISE research internship program, for his help with experiments. A. Cipitria would like to acknowledge funding from the DFG Emmy Noether grant (CI 203/2-1), from IKERBASQUE Basque Foundation for Science, and from the Spanish Ministry of Science and Innovation (PID2021-123013OB-I00).

## Supporting Information

Supplementary materials, including list of abbreviations used, details on methodology, theoretical derivations, and additional vesicle and hydrogel characterization data

## Conflict of Interest Disclosure

The authors declare no conflict of interest.

## Author Contributions

R. Dimova and A. Cipitria designed the project. N.W. Tam conducted experiments and wrote the manuscript. O. Schullian analyzed experimental data and provided theoretical derivations. R. Dimova and A. Cipitria supervised the project and edited the manuscript.

## Notes

### Competing Interest Statement

The authors have declared no competing interest.

### Summary of Updates

The key modifications include additional details about particle tracking analysis, new data illustrating diffusion in gels of varying agarose concentration, particle size distributions, fluorescence intensity analysis of aggregates and clarified the text on multiple locations.

## REFERENCES

1. Hessvik, N.P., and A. Llorente. 2018. Current knowledge on exosome biogenesis and release. Cell. Mol. Life Sci. 75:193–208.

2. Ono, M., N. Kosaka, N. Tominaga, Y. Yoshioka, F. Takeshita, R.-u. Takahashi, M. Yoshida, H. Tsuda, K. Tamura, and T. Ochiya. 2014. Exosomes from bone marrow mesenchymal stem cells contain a microRNA that promotes dormancy in metastatic breast cancer cells. Sci. Signal. 7:ra63–ra63.

3. Kooijmans, S.A.A., P. Vader, S.M. van Dommelen, W.W. van Solinge, and R.M. Schiffelers. 2012. Exosome mimetics: A novel class of drug delivery systems. Int. J. Nanomedicine. 7:1525–1541.

4. Chen, C.C., L. Liu, F. Ma, C.W. Wong, X.E. Guo, J. V. Chacko, H.P. Farhoodi, S.X. Zhang, J. Zimak, A. Ségaliny, M. Riazifar, V. Pham, M.A. Digman, E.J. Pone, and W. Zhao. 2016. Elucidation of Exosome Migration Across the Blood–Brain Barrier Model In Vitro. Cell. Mol. Bioeng. 9:509–529.

5. L. Arias, J. B. Clares, M. E. Morales, V. Gallardo, and M. A. Ruiz. 2011. Lipid-Based Drug Delivery Systems for Cancer Treatment. Curr. Drug Targets. 12:1151–1165.

6. Ozpolat, B., A.K. Sood, and G. Lopez-Berestein. 2014. Liposomal siRNA nanocarriers for cancer therapy. Adv. Drug Deliv. Rev. 66:110–6.

7. Sabanovic, B., F. Piva, M. Cecati, and M. Giulietti. 2021. Promising Extracellular Vesicle-Based Vaccines against Viruses, Including SARS-CoV-2. Biology (Basel). 10:94.

8. Lieleg, O., R.M. Baumgärtel, and A.R. Bausch. 2009. Selective filtering of particles by the extracellular matrix: An electrostatic bandpass. Biophys. J. 97:1569–1577.

9. Yu, M., W. Song, F. Tian, Z. Dai, Q. Zhu, E. Ahmad, S. Guo, C. Zhu, H. Zhong, Y. Yuan, T. Zhang, X. Yi, X. Shi, Y. Gan, and H. Gao. 2019. Temperature- and rigidity-mediated rapid transport of lipid nanovesicles in hydrogels. Proc. Natl. Acad. Sci. 116:5362–5369.

10. Lenzini, S., R. Bargi, G. Chung, and J.-W. Shin. 2020. Matrix mechanics and water permeation regulate extracellular vesicle transport. Nat. Nanotechnol. 15:217–223.

11. Zhang, X., J. Hansing, R.R. Netz, and J.E. DeRouchey. 2015. Particle Transport through Hydrogels Is Charge Asymmetric. Biophys. J. 108:530–539.

12. Xue, C., X. Shi, Y. Tian, X. Zheng, and G. Hu. 2020. Diffusion of Nanoparticles with Activated Hopping in Crowded Polymer Solutions. Nano Lett. 20:3895–3904.

13. Xu, Q., L.M. Ensign, N.J. Boylan, A. Schön, X. Gong, J.-C. Yang, N.W. Lamb, S. Cai, T. Yu, E. Freire, and J. Hanes. 2015. Impact of Surface Polyethylene Glycol (PEG) Density on Biodegradable Nanoparticle Transport in Mucus ex Vivo and Distribution in Vivo. ACS Nano. 9:9217–9227.

14. Xue, C., Y. Huang, X. Zheng, and G. Hu. 2022. Hopping Behavior Mediates the Anomalous Confined Diffusion of Nanoparticles in Porous Hydrogels. J. Phys. Chem. Lett. 13:10612–10620.

15. Burla, F., T. Sentjabrskaja, G. Pletikapic, J. Van Beugen, and G.H. Koenderink. 2020. Particle diffusion in extracellular hydrogels. Soft Matter. 16:1366–1376.

16. Jiang, L., and S. Granick. 2017. Real-Space, in Situ Maps of Hydrogel Pores. ACS Nano. 11:204–212.

17. Rodríguez-Suárez, J.M., C.S. Butler, A. Gershenson, and B.L.T. Lau. 2020. Heterogeneous Diffusion of Polystyrene Nanoparticles through an Alginate Matrix: The Role of Cross-linking and Particle Size. Environ. Sci. Technol. 54:5159–5166.

18. Yu, S., F. Tian, and X. Shi. 2022. Diffusion of deformable nanoparticles in adhesive polymeric gels. J. Mech. Phys. Solids. 167:105002.

19. Du, H., P. Chandaroy, and S.W. Hui. 1997. Grafted poly-(ethylene glycol) on lipid surfaces inhibits protein adsorption and cell adhesion. Biochim. Biophys. Acta - Biomembr. 1326:236–248.

20. Lee, H., and R.G. Larson. 2016. Adsorption of Plasma Proteins onto PEGylated Lipid Bilayers: The Effect of PEG Size and Grafting Density. Biomacromolecules. 17:1757–1765.

21. Garbuzenko, O., Y. Barenholz, and A. Priev. 2005. Effect of grafted PEG on liposome size and on compressibility and packing of lipid bilayer. Chem. Phys. Lipids. 135:117–129.

22. Jokerst, J. V., T. Lobovkina, R.N. Zare, and S.S. Gambhir. 2011. Nanoparticle PEGylation for imaging and therapy. Nanomedicine. 6:715–728.

23. Kaufmann, S., O. Borisov, M. Textor, and E. Reimhult. 2011. Mechanical properties of mushroom and brush poly(ethylene glycol)-phospholipid membranes. Soft Matter. 7:9267.

24. Wiklander, O.P.B., J.Z. Nordin, A. O’Loughlin, Y. Gustafsson, G. Corso, I. Mäger, P. Vader, Y. Lee, H. Sork, Y. Seow, N. Heldring, L. Alvarez-Erviti, C.E. Smith, K. Le Blanc, P. Macchiarini, P. Jungebluth, M.J.A. Wood, and S. EL Andaloussi. 2015. Extracellular vesicle in vivo biodistribution is determined by cell source, route of administration and targeting. J. Extracell. Vesicles. 4:26316.

25. Sbalzarini, I.F., and P. Koumoutsakos. 2005. Feature point tracking and trajectory analysis for video imaging in cell biology. J. Struct. Biol. 151:182–195.

26. Stellwagen, J., and N.C. Stellwagen. 1995. Internal Structure of the Agarose Gel Matrix. J. Phys. Chem. 99:4247–4251.

27. Narayanan, J., J.Y. Xiong, and X.Y. Liu. 2006. Determination of agarose gel pore size: Absorbance measurements vis a vis other techniques. J. Phys. Conf. Ser. 28:83–86.

28. Zarrintaj, P., S. Manouchehri, Z. Ahmadi, M.R. Saeb, A.M. Urbanska, D.L. Kaplan, and M. Mozafari. 2018. Agarose-based biomaterials for tissue engineering. Carbohydr. Polym. 187:66–84.

29. Aymard, P., D.R. Martin, K. Plucknett, T.J. Foster, A.H. Clark, and I.T. Norton. 2001. Influence of thermal history on the structural and mechanical properties of agarose gels. Biopolymers. 59:131–144.

30. Normand, V., D.L. Lootens, E. Amici, K.P. Plucknett, and P. Aymard. 2000. New Insight into Agarose Gel Mechanical Properties. Biomacromolecules. 1:730–738.

31. Gasperini, L., J.F. Mano, and R.L. Reis. 2014. Natural polymers for the microencapsulation of cells. J. R. Soc. Interface. 11:20140817.

32. Lee, K.Y., and D.J. Mooney. 2001. Hydrogels for Tissue Engineering. Chem. Rev. 101:1869–1880.

33. Thomsen, A.R., C. Aldrian, P. Bronsert, Y. Thomann, N. Nanko, N. Melin, G. Rücker, M. Follo, A.L. Grosu, G. Niedermann, P.G. Layer, A. Heselich, and P.G. Lund. 2018. A deep conical agarose microwell array for adhesion independent three-dimensional cell culture and dynamic volume measurement. Lab Chip. 18:179–189.

34. Mercey, E., P. Obeïd, D. Glaise, M.-L. Calvo-Muñoz, C. Guguen-Guillouzo, and B. Fouqué. 2010. The application of 3D micropatterning of agarose substrate for cell culture and in situ comet assays. Biomaterials. 31:3156–3165.

35. Cambria, E., S. Brunner, S. Heusser, P. Fisch, W. Hitzl, S.J. Ferguson, and K. Wuertz-Kozak. 2020. Cell-Laden Agarose-Collagen Composite Hydrogels for Mechanotransduction Studies. Front. Bioeng. Biotechnol. 8.

36. Lewitus, D.Y., J. Landers, J.R. Branch, K.L. Smith, G. Callegari, J. Kohn, and A. V. Neimark. 2011. Biohybrid Carbon Nanotube/Agarose Fibers for Neural Tissue Engineering. Adv. Funct. Mater. 21:2624–2632.

37. López-Marcial, G.R., A.Y. Zeng, C. Osuna, J. Dennis, J.M. García, and G.D. O’Connell. 2018. Agarose-Based Hydrogels as Suitable Bioprinting Materials for Tissue Engineering. ACS Biomater. Sci. Eng. 4:3610–3616.

38. Schwendener, R.A. 2014. Liposomes as vaccine delivery systems: a review of the recent advances. Ther. Adv. Vaccines. 2:159–182.

39. Faizi, H.A., C.J. Reeves, V.N. Georgiev, P.M. Vlahovska, and R. Dimova. 2020. Fluctuation spectroscopy of giant unilamellar vesicles using confocal and phase contrast microscopy. Soft Matter. 16:8996–9001.

40. Gracià, R.S., N. Bezlyepkina, R.L. Knorr, R. Lipowsky, and R. Dimova. 2010. Effect of cholesterol on the rigidity of saturated and unsaturated membranes: fluctuation and electrodeformation analysis of giant vesicles. Soft Matter. 6:1472.

41. Weinberger, A., F.-C. Tsai, G.H. Koenderink, T.F. Schmidt, R. Itri, W. Meier, T. Schmatko, A. Schröder, and C. Marques. 2013. Gel-Assisted Formation of Giant Unilamellar Vesicles. Biophys. J. 105:154–164.

42. Dimova, R., and C. Marques. 2020. The Giant Vesicle Book. Taylor & Francis.

43. Michalet, X. 2010. Mean square displacement analysis of single-particle trajectories with localization error: Brownian motion in an isotropic medium. Phys. Rev. E. 82:041914.

44. Wolde-Kidan, A., A. Herrmann, A. Prause, M. Gradzielski, R. Haag, S. Block, and R.R. Netz. 2021. Particle Diffusivity and Free-Energy Profiles in Hydrogels from Time-Resolved Penetration Data. Biophysj. 120:463–475.

45. Arends, F., R. Baumgärtel, and O. Lieleg. 2013. Ion-Specific Effects Modulate the Diffusive Mobility of Colloids in an Extracellular Matrix Gel. Langmuir. 29:15965–15973.

46. Faizi, H.A., S.L. Frey, J. Steinkühler, R. Dimova, and P.M. Vlahovska. 2019. Bending rigidity of charged lipid bilayer membranes. Soft Matter. 15:6006–6013.

47. Li, S.-J., and X. Shi. 2021. Tailoring Antifouling Properties of Nanocarriers via Entropic Collision of Polymer Grafting. ACS Nano. 15:5725–5734.

48. Israelachvili, J.N. 2011. Intermolecular and Surface Forces. 3rd ed. Burlington: Elsevier.

49. Marsh, D., R. Bartucci, and L. Sportelli. 2003. Lipid membranes with grafted polymers: physicochemical aspects. Biochim. Biophys. Acta - Biomembr. 1615:33–59.

50. Hiergeist, C., and R. Lipowsky. 1996. Elastic Properties of Polymer-Decorated Membranes. J. Phys. II. 6:1465–1481.

51. Mertins, O., and R. Dimova. 2013. Insights on the Interactions of Chitosan with Phospholipid Vesicles. Part II: Membrane Stiffening and Pore Formation. Langmuir. 29:14552–14559.

52. Gompper, G., H. Endo, M. Mihailescu, J. Allgaier, M. Monkenbusch, D. Richter, B. Jakobs, T. Sottmann, and R. Strey. 2001. Measuring bending rigidity and spatial renormalization in bicontinuous microemulsions. Europhys. Lett. 56:683–689.

