## Supporting information for "Nonspecific Membrane-Matrix Interactions Influence Diffusivity of Lipid Vesicles in Hydrogels"

### Supplementary Materials

Nicky W. Tam,[1] Otto Schullian,[1,2] Amaia Cipitria,[1,3,4] and Rumiana Dimova[1]*

[1]Max Planck Institute of Colloids and Interfaces, Science Park Golm, 14476 Potsdam, Germany
[2]Free University of Berlin, Department of Physics, 14195 Berlin, Germany
[3]Biodonostia Health Research Institute, San Sebastián, Spain
[4]IKERBASQUE, Basque Foundation for Science, Bilbao, Spain

**List of abbreviations used**

| | |
|---|---|
| **DiI** | 1,1'-Dioctadecyl-3,3,3',3'-Tetramethylindodicarbocyanine (lipophilic dye) |
| **DOPC** | 1,2-dioleoyl-sn-glycero-3-phosphocholine (neutral phospholipid) |
| **DOPS** | 1,2-dioleoyl-sn-glycero-3-phospho-L-serine (negatively charged phospholipid) |
| **DOTAP** | 1,2-dioleoyl-3-trimethylammonium-propane (positively charged surfactant) |
| **DSPE-mPEG1K, -2K, -5K** | 1,2-distearoyl-sn-glycero-3-phosphoethanolamine-N-[methoxy(polyethylene glycol)-1000], -2000], and -5000] (PEGylated lipids) |
| **ECM** | Extracellular matrix |
| **EV** | Extracellular vesicle |
| **GUV** | Giant unilamellar vesicle |
| **LUV** | Large unilamellar vesicle |
| **MSD** | Mean squared displacement |
| **MW** | Molecular weight |
| **PBS** | Phosphate buffered saline |
| **PEG** | Polyethylene glycol |
| **PVA** | Polyvinyl alcohol |
| **ROI** | Region of interest |
| **WLE** | Wavelength exponent |

### Section S1. PVA-assisted swelling of GUVs and their bending rigidity

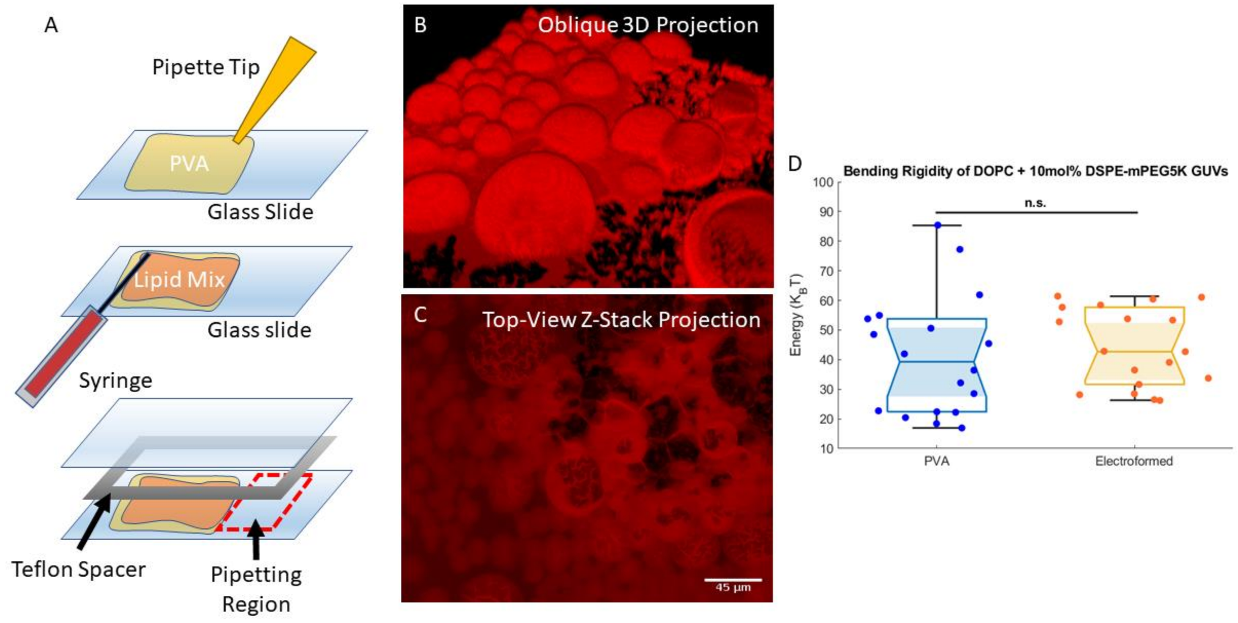


*Figure S1, PVA gel-assisted swelling of giant unilamellar vesicles (GUVs).* (A) Schematic presentation of the approach. A layer of 5% w/v polyvinyl alcohol (PVA) is spread onto a glass slide with a pipette tip and dried. Next, a solution of lipids dissolved in chloroform is spread on top with a syringe and dried under vacuum. The swelling chamber is assembled by sandwiching a Teflon spacer between the coated slide and a second clean slide. Buffer is added to the chamber and the lipid film is left to swell over 30 minutes, resulting in the formation of GUVs. The GUVs are afterwards harvested by pipetting solution from a PVA- and lipid-free region of the chamber. (B,C) Three-dimensional confocal projection (B) and a top-down Z-stack (C) of GUVs composed of DOPC forming on a layer of PVA (the same area is imaged in both panels). The lipid bilayer is visualized with a lipophilic dye, DiI. (D) Comparison of bending rigidities between DOPC + 10mol% DSPE-mPEG5K GUVs formed with PVA-assisted swelling (PVA) and standard electroformation (Electroformed), as measured with fluctuation analysis. While PVA-swelled GUVs have greater variance in bending rigidity values, they have a similar mean value comparable to electroformed GUVs. A paired-sample t-test did not find any statistically significant difference in values ($p>0.05$).

The PVA-assisted swelling of GUVs follows the protocol reported by Weinberger et al.(1) First, 20 µL 5% w/v solution of polyvinyl alcohol (PVA; fully hydrolyzed, MW = 145000 Da; Merck Group) in water with 50 mM sucrose was spread onto a 2 cm by 5 cm glass slide area, slightly smaller than the dimensions of a rectangular, 2 mm-thick Teflon spacer (see Fig. S1A), and allowed to dry completely in an oven at 50°C. Next, a thin 15 µL layer of 4 mM lipid mixture dissolved in chloroform was spread on top of the PVA layer and dried in a vacuum for 1.5 hours. The slide was then assembled into a sandwich with another glass slide and a Teflon spacer in the middle, held together with binder clips (Fig. S1). The lipid layer was hydrated for 30 minutes with 2 mL PBS + 50 mM sucrose (345 mOsm/kg). The sucrose was necessary to help with the swelling process and to generate a sugar gradient that would later aid in visualizing the GUVs. The resulting osmolality of the GUV solution was 358 mOsm/kg, as measured with an Osmomat 3000 freezing point osmometer (Gonotech), suggesting only partial recovery of the sucrose from the PVA layer. GUVs were harvested very carefully by pipetting liquids over a small region of the glass slide on one end of the spacer that was left bare without any PVA or lipid. They were then diluted 1:1 in a solution of PBS + 100mM glucose (394mOsm/kg) to slightly deflate the GUVs, which was necessary for visualization in phase contrast and to obtain the thermal membrane fluctuations required for fluctuation analysis. For the fluctuation analysis, we carefully selected the vesicles to be measured, avoiding GUVs that appear denser under phase contrast, which might suggest the presence of PVA in their lumen.

We used PVA-assisted swelling due to the poor compatibility of standard electroformation protocols with high salt content buffers. We wanted to recapitulate physiological salt concentrations for our experiments with LUVs, and thus we aimed to recreate those conditions with our GUVs for fluctuation analysis. PVA-assisted swelling can result in PVA contamination in the resulting GUVs, thus altering the stretching elasticity modulus of the membrane (2). However, this did not appear to greatly affect the measured bending rigidity when compared to GUVs formed with standard electroformation, similarly to previous data on single-component lipid membranes (3). For electroformed GUVs, a thin layer of 10 μL lipids dissolved in chloroform was spread onto the conductive surfaces of two indium-tin-oxide- (ITO) coated glass plates. This was dried under vacuum for 1 hour before being assembled into a swelling chamber with a 2mm-thick Teflon spacer, similar to the chamber sketched in Fig. S1A. The lipid layer was rehydrated with a 20 mM sucrose solution and a 10Hz, 1.6$V_{pp}$ AC electric field supplied by a function generator was passed across the glass plates for 1 hour to induce GUV formation. GUVs were harvested by pipetting and diluted 1:10 in a solution of 22mM glucose prior to imaging for fluctuation analysis. Fig. S1D shows that while PVA-swelled GUVs have higher variance in bending rigidity than electroformed GUVs, possibly due to PVA contamination, both populations of GUVs have a similar mean bending rigidity.

#### Section S2. Turbidimetric assay for average agarose gel pore size

Molten agarose in distilled water was added to standard 1cm PMMA cuvettes and allowed to cool to room temperature (~22°C) over 10 minutes to gel. Absorbance values, $\alpha$, over a range of wavelengths, λ, 600-900nm were measured and a blank signal obtained from a cuvette filled with distilled water was removed by subtraction. These were then converted to turbidity, τ, with the following formula(4)

$$\tau(\lambda) = \frac{2.3\alpha(\lambda)}{L} \qquad (1)$$

where $L$ is the optical path length. Linear regression was done on a double-log plot of the turbidity as a function of the wavelength to obtain the wavelength exponent, WLE= $\frac{\mathrm{d}\log\tau(\lambda)}{\mathrm{d}\log\alpha(\lambda)}$. This WLE was then compared to data derived numerically from the analytical results of Aymard et al.(5) relating WLE and the correlation length; the latter can be roughly assumed equal to the pore size, see Fig. S4.

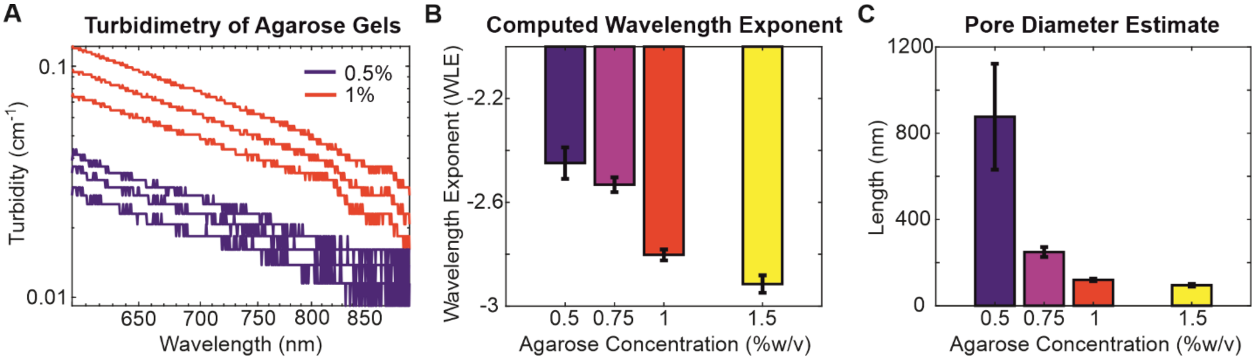


*Figure S2, Estimation of average agarose gel pore size by turbidimetry.* (A) Representative turbidity data for 0.5% and 1% agarose gels, used to determine wavelength exponent (WLE). Each individual curve represents a separate sample with total n=3 for each concentration. (B) WLE values obtained for agarose gels of different concentrations. (C) Average pore size estimated using analytical data from Aymard et al.(5) The high variability in the 0.5% agarose gel arises from the fact that this concentration lies at the lower limit for which this method is valid.

### Section S3. Mobility cutoff and mobile fraction

To form 1% agarose gels with embedded LUVs, 25µL 2% agarose stock solution was mixed with 24µL PBS and 1µL LUV solution directly on a glass microscope slide with a rubber spacer set on a hotplate at 35°C by pipetting up and down (Fig. S3A). For 0.5% w/v agarose gels, the amount of agarose stock used was reduced to 12.5µL and the PBS raised to 36.5µL, keeping the LUV solution at 1µL. A list of gel formulations can be found in Table 2 in Section S5.

As explained in the main text, the mobile fraction, or the proportion of particles that are not immobilized in the gel is represented by the sum of the bins of the histograms of log diffusion coefficients greater than the mobility cutoff of -14. Experimentally, the histograms of these particle populations appear to have a strong bimodal distribution, especially in conditions where there is a large proportion of mobile particles, such as in 0.5% agarose. In such cases, the peaks of the bimodal distributions are well-separated by the mobility cutoff of -14 (see example histograms in Fig. S3C,D).

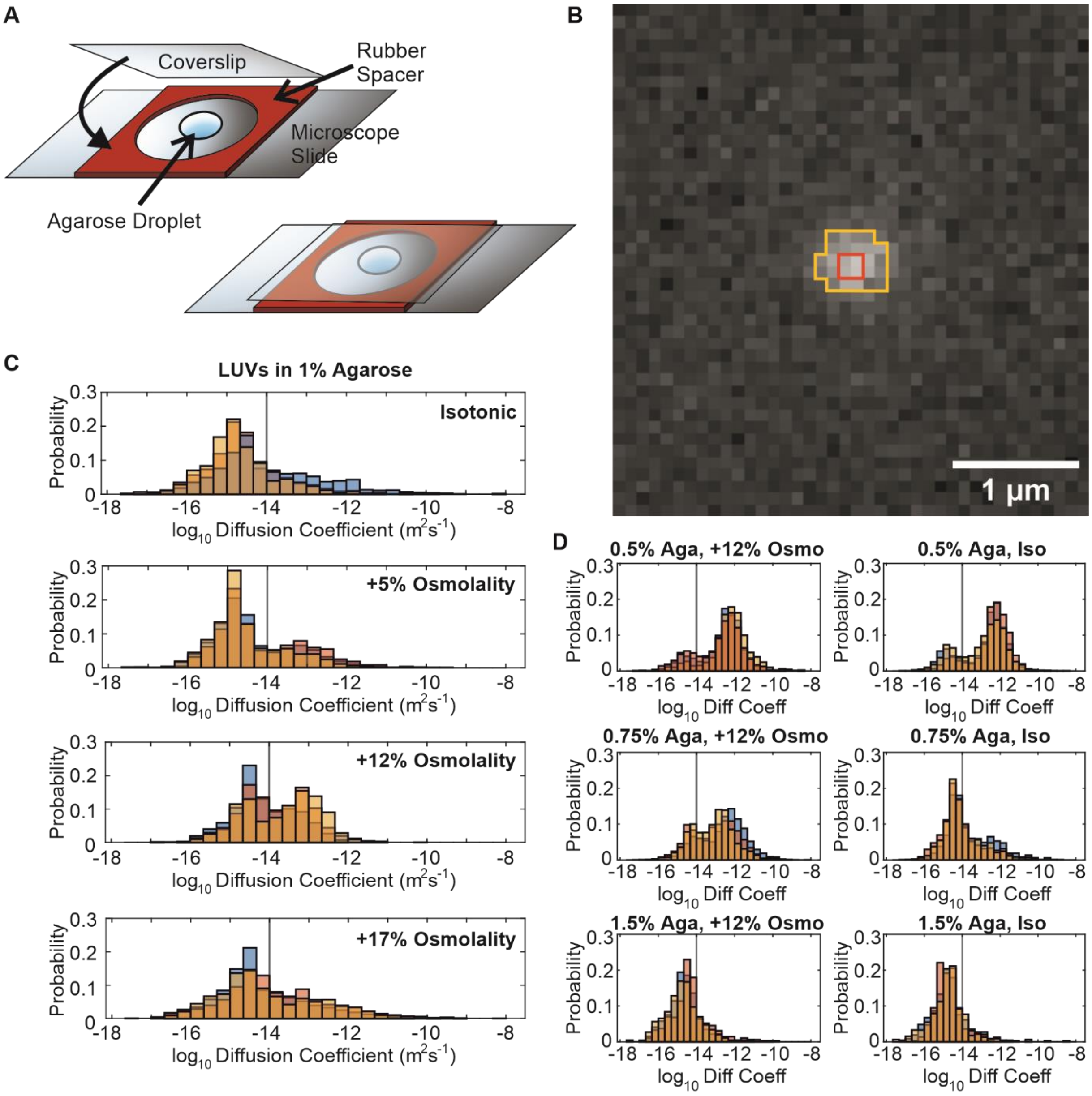


*Figure S3, Single particle tracking of DOPC LUVs in agarose gels with varying osmolality.* (A) General experimental setup for single particle tracking experiments on embedded DOPC LUVs. Agarose gels and LUVs are mixed directly on a confocal slide with a rubber spacer and then sealed with a coverslip placed on top to make a gel disc. (B) Representative image of a ~100nm DOPC LUV fluorescently labelled with DiI. Due to being below the diffraction limit, the particle appears as a cluster of 4 bright pixels (highlighted in red) surrounded by 1-2 pixel spread (highlighted in yellow). (C) Histograms of $\log_{10}$ diffusion coefficients of DOPC LUVs in 1% agarose gels of different osmotic conditions. Each colour represents a different replicate, with pooled data from particles in three different regions of interest per replicate. The mobility cutoff of -14 is indicated with a vertical line, showing the separation of mobile and immobile regions of the distributions. (D) Histograms of $\log_{10}$ diffusion coefficients of DOPC LUVs in 0.5%, 0.75%, and 1.5% agarose gels with different osmotic conditions. Only the +12% osmolality condition was tested along with the isotonic condition because it showed the greatest difference in mobile fraction at 1% agarose concentration.

### Section S4. Derivation of a collective diffusion coefficient from infiltrating LUVs

The prior imaging chamber setup was modified to study the infiltration of LUVs into preformed agarose disks. First, the rubber spacer was sealed with vacuum grease to the glass slide to prevent evaporation over 100 hours of imaging. With the imaging chamber open and sitting on a hotplate at 35°C, a 1% w/v agarose gel was mixed without LUV solution directly on the slide by pipetting up and down. Next, a coverslip was placed on top of the liquid gel droplet, such that the droplet wetted both glass surfaces and formed a disk, but leaving a small opening between the spacer and the coverslip for introducing the LUV solution later, see Fig. S4A. The imaging chamber was taken off the hotplate to cool at room temperature for 5 minutes. Once the gel had set, a solution of LUVs diluted 1:3 after extrusion (to a final lipid concentration of 12 μM) was pipetted into the imaging chamber via the opening, where it was wicked towards the gel. The chamber was then sealed with more vacuum grease to cover the opening.

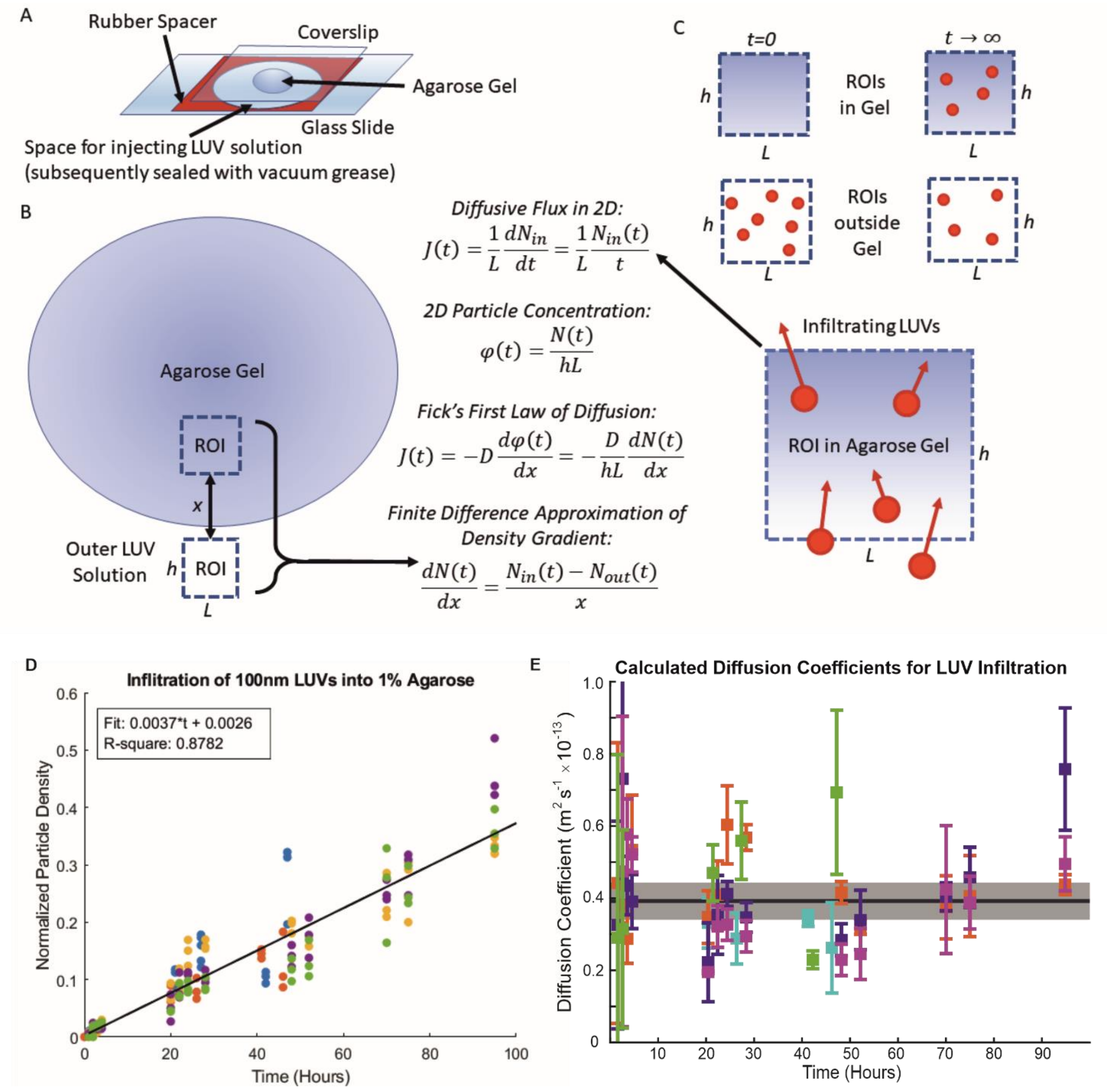


*Figure S4, Infiltration of DOPC LUVs into agarose gels and the derivation of the diffusion coefficient based on population dynamics.* A) A sketch of the imaging chamber for preparing the agarose gel disk (sandwiched between the glass slide and cover slip) around which the DOPC LUV solution is introduced, followed by monitoring of LUV infiltration. B) Schematic of the sample with relative positions of the regions of interest (ROIs). The dimensions of the ROI are $h$ and $L$, and the ROIs inside and outside of the gel are separated by a distance, $x$ (typically 300 μm). The entire gel disc has an approximate diameter of 8mm. C) Schematic diagram of the ROIs and the location of LUVs within them at different times. At time, $t$ = 0, all LUVs are outside of the gel as none of them have infiltrated yet. Over time, LUVs infiltrate into the gel, whose diffusive flux, $J(t)$ can be calculated as shown; see text for details. We note that equilibrium was never reached in the 100-hour experimental timeframe. D) Normalized particle density in the gel interior as a function of time. Particles within the interior ROI are counted and normalized by the number of particles in the corresponding exterior ROI for each time point. Different colours represent different experimental replicates. Particle density appears to increase linearly with time and is presented here with a linear least-squares fit. E) Diffusion coefficients determined at each time point. Different colours represent different replicates. Error bars show standard deviation across three ROIs at each timepoint, with the black line and gray shaded area showing the ensemble average and standard deviation across replicates. The data is also shown in Table 1.

Images of the gel interior and of the external LUV solution were taken at three regions of interest (ROIs) per gel over 100 hours. ROIs in the gel interior were chosen approximately 300μm from the edge of the gel (see schematic in Fig. S4B) to avoid the crowded LUVs skewing the dynamic range of the camera. Vertical position in the gel was held approximately constant by identifying the position of the surface of the glass slide, at which adhered LUVs could be identified and moving up the sample by a full turn of the focus knob. The particle flux did not reach equilibrium over 100 hours of monitoring, as the diffusion was very slow.

The results of five independent replicates are shown in Table 1 and give an ensemble average of $D = 3.92 \pm 0.52 \times 10^{-14}$ m$^2$ s$^{-1}$, corresponding to a $\log_{10}$ value of -13.41.

| Measurement No. | $D \times 10^{-14} \mathrm{m}^2\mathrm{s}^{-1}$ |
|---|---|
| 1 | $3.08 \pm 0.38$ |
| 2 | $4.26 \pm 0.94$ |
| 3 | $4.21 \pm 1.57$ |
| 4 | $3.76 \pm 1.24$ |
| 5 | $4.28 \pm 1.79$ |
| Average | $3.92 \pm 0.52$ |

*Table 1,* Diffusion coefficients determined for the infiltration of LUVs into 1% agarose gels.

**Section S5. Osmotic effects on gels and LUVs**

For hypertonic agarose gels, 2% w/v agarose stocks made in PBS (final osmolality ~290mOsm/kg) were mixed in varying proportions with PBS and a stock solution of 100mM glucose dissolved in PBS (~390mOsm/kg). At 0.5% agarose concentration, only the 12% hypertonic condition was tested as it showed the greatest increase in LUV mobility in 1% agarose gels (see histograms in Fig. S3).

| Final Agarose Concentration | Final Gel Volume (μL) | Vol. 2% Agarose Stock (μL) | Vol. PBS diluent (μL) | Vol. PBS + glucose solution (μL) | Vol. LUV solution (μL) | Absolute Osmolality (mOsm/kg) | Relative Osmolality Change |
|---|---|---|---|---|---|---|---|
| 1.5% | 50 | 37 | 12 | 0 | 1 | 289 | 0 |
| 1.5% | 50 | 37 | 0 | 12 | 1 | 325 | +12% |
| 1% | 50 | 25 | 24 | 0 | 1 | 289 | 0 |
| 1% | 50 | 25 | 17 | 7 | 1 | 303 | +5% |
| 1% | 50 | 25 | 7 | 17 | 1 | 325 | +12% |
| 1% | 50 | 25 | 0 | 24 | 1 | 338 | +17% |
| 0.75% | 50 | 18.5 | 30.5 | 0 | 1 | 289 | 0 |
| 0.75% | 50 | 18.5 | 13.5 | 17 | 1 | 325 | +12% |
| 0.5% | 50 | 12.5 | 36.5 | 0 | 1 | 289 | 0 |
| 0.5% | 50 | 12.5 | 19.5 | 17 | 1 | 325 | +12% |

*Table 2*, Gel formulations for testing the osmotic effects on the diffusion coefficient of embedded LUVs in agarose gels.

Hypertonic agarose gels (320mOsm/kg) relative to intravesicular solutions used in experiments (290mOsm/kg) were formed by addition of glucose to molten agarose solutions prior to gelation. This was done to determine whether the presence of glucose would affect agarose gel mechanics (Fig. S5A). No significant differences were observed in 0.5% and 1% agarose gels.

Osmotic effects on LUVs were measured by DLS independently of samples used for embedding in gels (Fig. S5B). LUVs were extruded with initial intravesicular solutions of 290mOsm/kg and were subsequently incubated in hypotonic (270mOsm/kg) and hypertonic solutions (320mOsm/kg and 390mOsm/kg). This was done for LUVs extruded through 100nm and 200nm pore size polycarbonate membranes to determine whether any osmotic effects would be more pronounced in larger LUVs. No significant differences in sizes due to osmolality were observed. Differences in sizes at the levels of hypertonic deflation achieved with LUVs embedded in gels and in the independent measurements with 100nm and 200nm LUVs under different osmotic conditions would not be readily measurable with DLS. This is because an osmotic difference of 12% resulting in a volume reduction of 12% corresponds to only 4% decrease in liposome diameter, which is at the detection limit of the device. We note that DOPC LUVs extruded through a 100nm-sized polycarbonate membrane appear as particles with 150nm diameter in DLS measurements. In the main text, we refer to these as 100nm-extruded LUVs because DLS measurements can be affected by membrane fluctuations and to emphasize that they are still capable of passing through 100nm-sized pores, such as those in the extruder membrane used to make them.

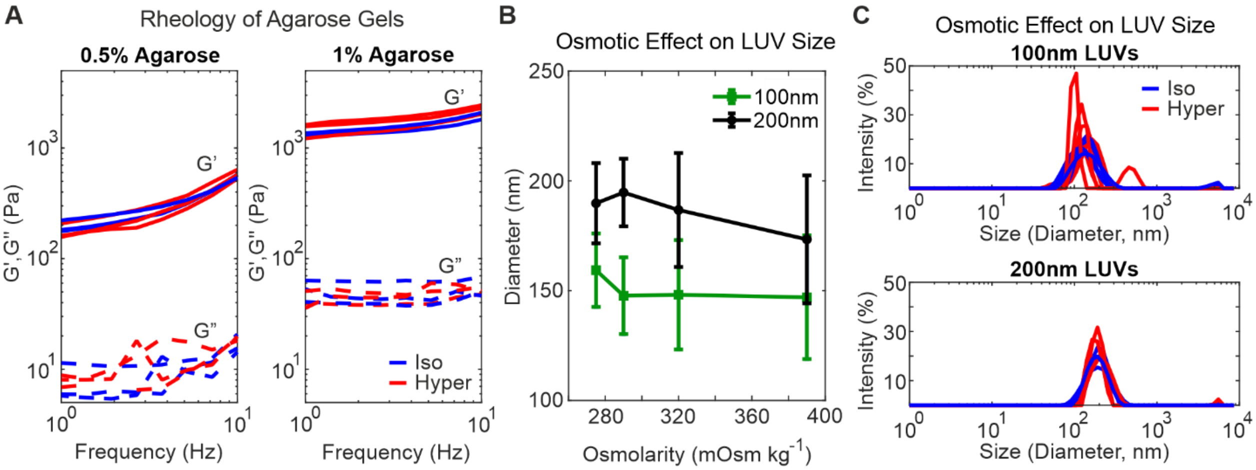


*Figure S5, Osmotic effects on gels and LUVs.* A) Bulk rheology of 0.5% and 1% agarose gels made in isotonic (290mOsm/kg, blue) and hypertonic (320mOsm/kg, red) solutions, relative to initial LUV interior solutions used in embedded LUV experiments. Neither storage (G') nor loss (G'') moduli appear to change with osmolality conditions. B) Average sizes of LUVs (with initial internal osmolality of 290mOsm/kg) when exposed to solutions of different osmolalities, as measured with DLS. The first data point in each series corresponds to a hypotonic condition, the second to isotonic condition, and the final two to hypertonic conditions. Different extruder pore sizes (100nm – green, and 200nm - black) were used to make LUVs to determine if osmotic inflation or deflation had a more pronounced effect on larger LUVs, but none was observed. Error bars show standard deviation. C) Representative size distributions of LUVs in osmotic conditions corresponding to those used in experiments with LUVs embedded in agarose, as measured with DLS. More variability can be seen in the LUVs exposed to 12% hypertonic (red; 320mOsm/kg) solution relative to interior osmolality compared to those exposed to isotonic solution (blue; 290mOsm/kg), likely due to membrane fluctuations.

### Section S6. Particles of varying surface charge and bulk mechanics

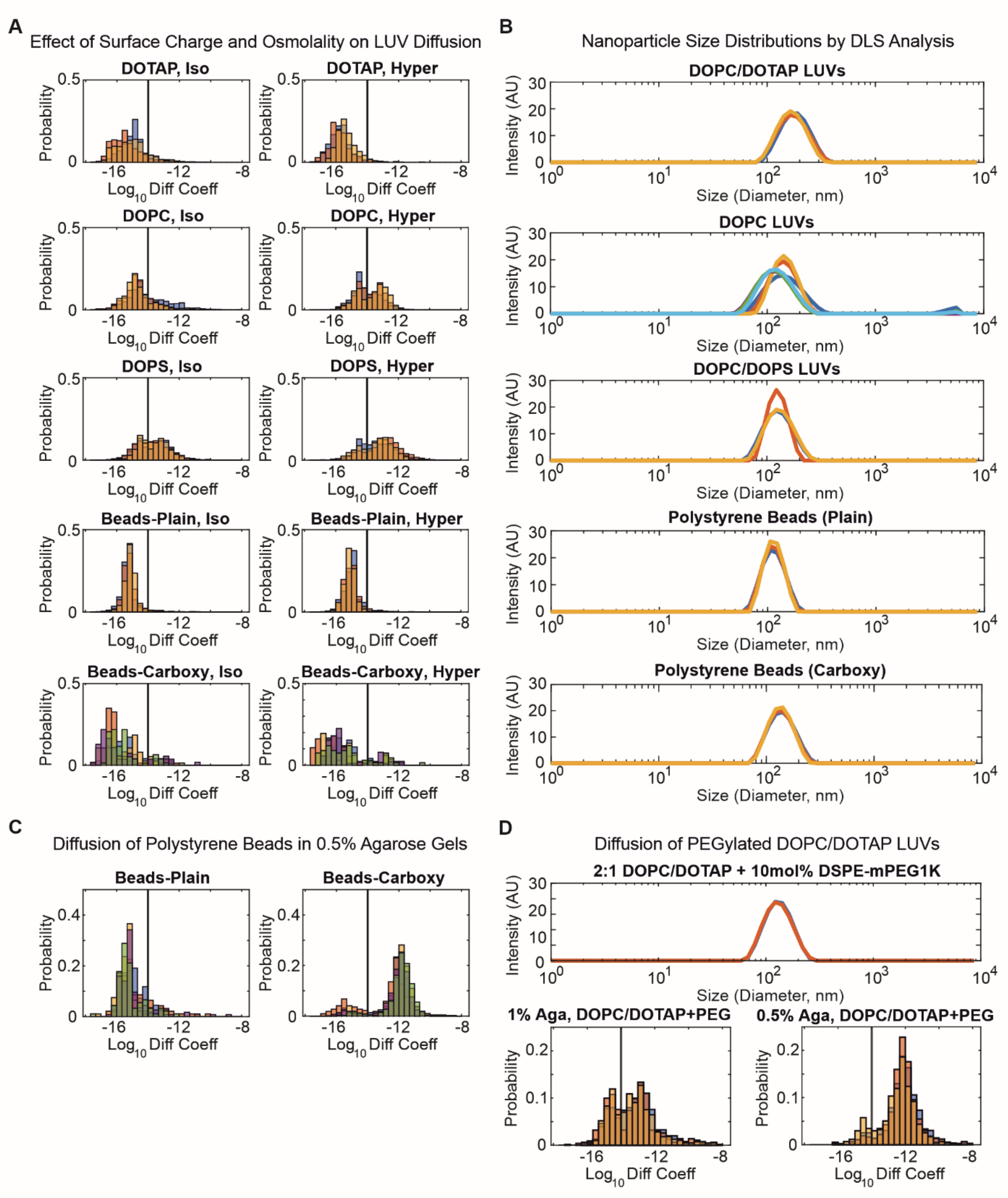


*Figure S6, Effect of charge and composition on particle mobility in agarose gels.* A) Histograms of $\log_{10}$ diffusion coefficients of particles with different surface charges in 1% isotonic (Iso; 290mOsm/kg) and hypertonic (Hyper, 320mOsm/kg) agarose gels: pure DOPC, 2:1 DOPC/DOPS, 2:1 DOPC/DOTAP, and polystyrene beads with (Beads-Carboxy) and without (Beads-Plain) surface carboxylation. B) Representative size distributions of the particles in A, as determined with DLS analysis. Different colours represent different batches of LUVs or different replicates. C) Histograms of $\log_{10}$ diffusion coefficients of polystyrene beads with (Beads-Carboxy) and without (Beads-Plain) surface carboxylation embedded in isotonic 0.5% agarose gels. D) Size distributions of 2:1 DOPC/DOTAP + 10mol% DSPE-mPEG1K LUVs, as measured with DLS (top), with histograms showing the distribution of $\log_{10}$ diffusion coefficients in 0.5% and 1% agarose (bottom). Different colours represent different replicates.

### Section S7. Passivation of LUVs with PEG

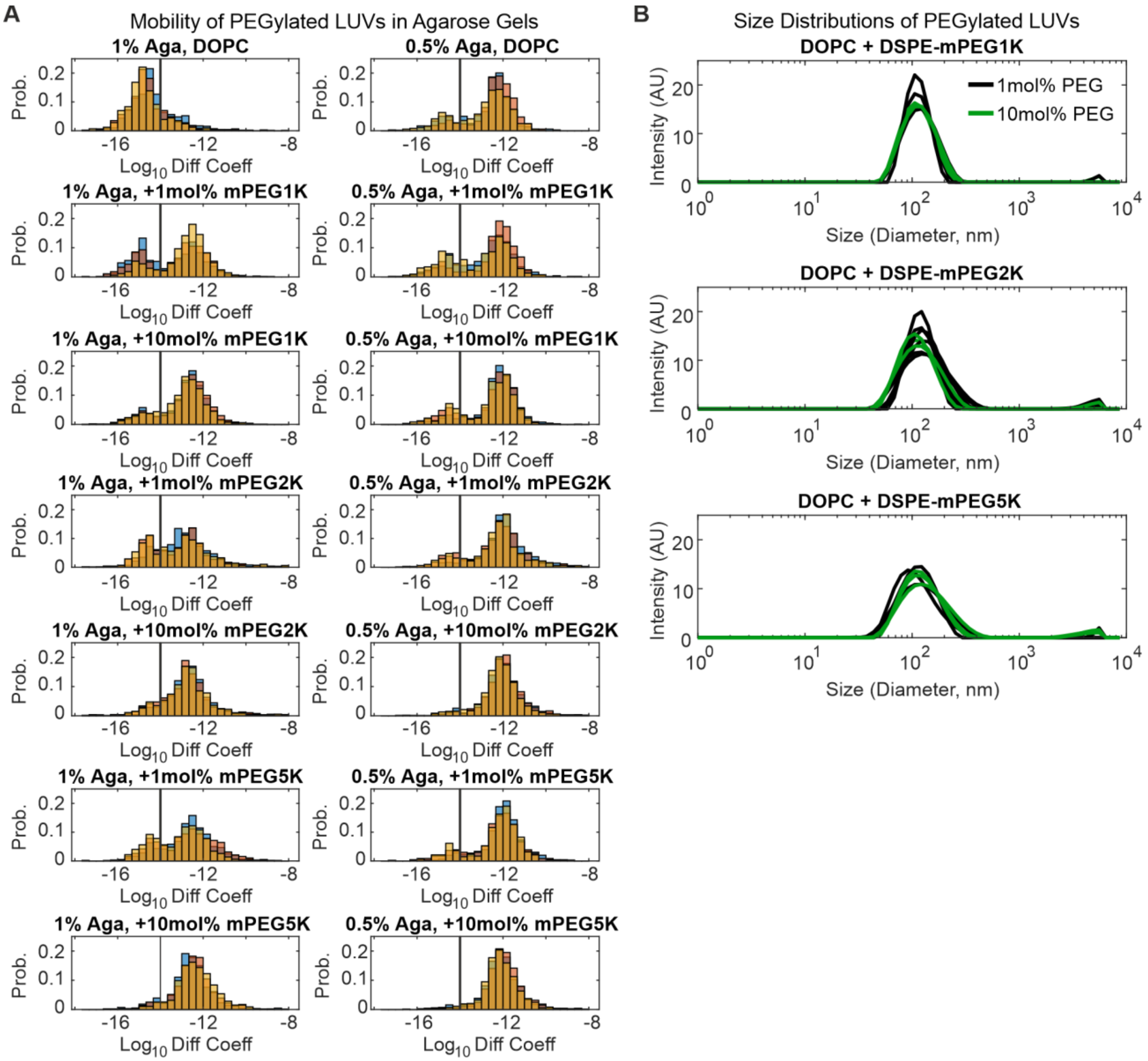


*Figure S7, Effect of PEGylation on LUV characteristics and mobility.* A) Histograms of $\log_{10}$ diffusion coefficients of DOPC LUVs doped with PEGylated lipids in 1% or 0.5% agarose gels (1% Aga, 0.5 % Aga). Different colours represent different replicates. The amount of PEGylated DSPE used to make the LUVs (in mol%) and the size of the PEG chain are indicated. B) Representative size distributions of LUVs made with PEGylated lipids of varying PEG chain size at 1mol% (black) or 10mol% (green) concentration in the membrane.

**References**


1. Weinberger, A., F.-C. Tsai, G.H. Koenderink, T.F. Schmidt, R. Itri, W. Meier, T. Schmatko, A. Schröder, and C. Marques. 2013. Gel-Assisted Formation of Giant Unilamellar Vesicles. *Biophys. J.* 105:154–164.

2. Dao, T.P.T., M. Fauquignon, F. Fernandes, E. Ibarboure, A. Vax, M. Prieto, and J.F. Le Meins. 2017. Membrane properties of giant polymer and lipid vesicles obtained by electroformation and pva gel-assisted hydration methods. *Colloids Surfaces A Physicochem. Eng. Asp.* 533:347–353.

3. Faizi, H.A., A. Tsui, R. Dimova, and P.M. Vlahovska. 2022. Bending Rigidity, Capacitance, and Shear Viscosity of Giant Vesicle Membranes Prepared by Spontaneous Swelling, Electroformation, Gel-Assisted, and Phase Transfer Methods: A Comparative Study. *Langmuir*. 38:10548–10557.

4. Narayanan, J., J.Y. Xiong, and X.Y. Liu. 2006. Determination of agarose gel pore size: Absorbance measurements vis a vis other techniques. *J. Phys. Conf. Ser.* 28:83–86.

5. Aymard, P., D.R. Martin, K. Plucknett, T.J. Foster, A.H. Clark, and I.T. Norton. 2001. Influence of thermal history on the structural and mechanical properties of agarose gels. *Biopolymers*. 59:131–144.
